# Tomato I2 immune receptor can be engineered to confer partial resistance to the oomycete *Phytophthora infestans* in addition to the fungus *Fusarium oxysporum*

**DOI:** 10.1101/022079

**Authors:** Artemis Giannakopoulou, John F. C. Steele, Maria Eugenia Segretin, Tolga O. Bozkurt, Ji Zhou, Silke Robatzek, Mark J. Banfield, Marina Pais, Sophien Kamoun

## Abstract

Plants and animals rely on immune receptors, known as nucleotide-binding domain and leucine-rich repeat containing proteins (NB-LRR or NLR), to defend against invading pathogens and activate immune responses. How NLR receptors respond to pathogens is inadequately understood. We previously reported single-residue mutations that expand the response of the potato immune receptor R3a to AVR3a^EM^, a stealthy effector from the late blight oomycete pathogen *Phytophthora infestans*. I2, another NLR that mediates resistance to the wilt causing fungus *Fusarium oxysporum f. sp. lycopersici*, is the tomato ortholog of R3a. We transferred previously identified R3a mutations to I2 to assess the degree to which the resulting I2 mutants have an altered response. We discovered that wild-type I2 protein responds weakly to AVR3a. One mutant in the N-terminal coiled-coil domain, I2^I141N^, appeared sensitized and displayed markedly increased response to AVR3a. Remarkably, I2^I141N^ conferred partial resistance to *P. infestans.* Further, I2^I141N^ has an expanded response spectrum to *F. oxysporum f. sp. lycopersici* effectors compared to the wild-type I2 protein. Our results suggest that synthetic immune receptors can be engineered to confer resistance to phylogenetically divergent pathogens and indicate that knowledge gathered for one NLR could be exploited to improve NLRs from other plant species.

## Introduction

Plant diseases are one of the main threats to modern human life, and the need for sustainable crop disease resistance is becoming increasingly urgent (Pennisi 2010, Fisher, Henk et al. 2012). Plants have evolved various disease resistance mechanisms to defend against pathogens. Sustainable crop improvement based on these mechanisms often involve *R* genes - plant loci that encode immune receptors (Cook 2000, Michelmore, Christopoulou et al. 2013, Jones, Witek et al. 2014). One challenge is to engineer plants with broad-spectrum disease resistance, i.e. that can defend against a wide spectrum of pathogens. Although this can be achieved by pyramiding multiple immune receptors with different pathogen resistance spectra, single *R* genes that function against multiple pathogens do occur. One example is the tomato gene *Mi-1.2*, which confers resistance to pathogens from different phyla: arthropods (aphids and whiteflies) and a nematode (Nombela, Williamson et al. 2003, Atamian, Eulgem et al. 2012). Cf-2, an extracellular tomato immune receptor also mediates resistance to both the fungus *Cladosporium fulvum* and the root parasitic nematode *Globodera rostochiensis* (Lozano-Torres, Wilbers et al. 2012). Moreover, in Arabidopsis, RPS4 and RRS1 mediate resistance to the bacteria *Pseudomonas syringae* and *Ralstonia solanacearum* but also to the fungus *Colletotrichum higginsianum* (Gassmann, Hinsch et al. 1999, Deslandes, Olivier et al. 2003, Birker, Heidrich et al. 2009, Narusaka, Shirasu et al. 2009). However, pathogens continuously evolve new races that overcome immunoreceptor-specific mediated disease resistance. The challenge for plant breeders and biotechnologists is to generate new resistance gene specificities rapidly enough to keep up with pathogen evolution. Engineering synthetic wide-spectrum immune receptors that target more than one pathogen is one approach to rapidly deliver agronomically useful resistance genes.

Plants resist pathogenic invasion through cell surface and intracellular immune receptors that recognize pathogen-secreted molecules and activate immune responses (Dodds and Rathjen 2010, Win, Chaparro-Garcia et al. 2012). The largest family of intracellular immune receptors is the nucleotide binding-leucine-rich repeat (NLR) protein family, an important element of defense against pathogens in both plants and animals (Maekawa, Kufer et al. 2011, Jacob, Vernaldi et al. 2013). In plants, NLR proteins recognize effectors, pathogen secreted molecules that normally modulate host cell processes to the benefit of the pathogen (Hogenhout, Van der Hoorn et al. 2009, Dodds and Rathjen 2010, Bozkurt, Schornack et al. 2012, Win, Chaparro-Garcia et al. 2012). NLRs tend to recognize one or a limited number of effectors, resulting in NLR-triggered immunity or effector-triggered immunity (Jones and Dangl 2006, Ooijen, Burg et al. 2007, Dodds and Rathjen 2010, Win, Chaparro-Garcia et al. 2012). There are two major classes of NLR proteins depending on their N terminal domains: CC-NB-LRR or CNL proteins with a predicted coiled-coil (CC) structure and TIR-NB-LRR or TNL proteins with a Toll/interleukin-1 receptor (TIR) domain (Pan, Wendel et al. 2000, Andolfo, Jupe et al. 2014). The nucleotide-binding (NB) domain, also known as NB-ARC (shared by the mammalian protein Apaf-1, plant resistance proteins, and the nematode *Caenorhabditis elegans* Ced-4 protein) is the most conserved among the NLR proteins and is thought to act as a switch between the resting (ADP-bound) and activated state (ATP-bound) of the receptor (Takken, Albrecht et al. 2006, Tameling, Vossen et al. 2006, Maekawa, Kufer et al. 2011, Takken and Goverse 2012). The C-terminal LRR domain tends to be more polymorphic with the number and length of the LRR repeats being highly variable between different NLR proteins (Takken and Goverse 2012). Activation of NLR receptors usually results in a localized cell death response known as the hypersensitive response (HR), which is associated with restricting pathogen colonization of the host tissue (Spoel and Dong 2012, Win, Chaparro-Garcia et al. 2012).

The CNL class of NLRs has dramatically expanded in the Solanaceae (nightshade) botanical family (Andolfo, Jupe et al. 2014). In tomato, for example, 18 distinct clades of CNL genes are present in every chromosome, whereas only a single TNL clade occurs (Andolfo, Jupe et al. 2014). Two agronomically important genes occur in the Solanaceae CNL-8 clade. *R3a* from the wild potato *Solanum demissum* confers resistance to specific strains of the late blight oomycete pathogen *Phytophthora infestans* that carry the avirulence effector AVR3a^KI^ (Armstrong, Whisson et al. 2005, Bos, Kanneganti et al. 2006). In tomato, the *R3a* ortholog *I2* confers resistance against race 2 of the ascomycete fungus *Fusarium oxysporum f. sp. lycopersici*, the agent of tomato wilt disease (Ori, Eshed et al. 1997, Simons, Groenendijk et al. 1998, Takken and Rep 2010). *F. oxysporum f. sp. lycopersici* secretes the effector AVR2, which activates I2-mediated immunity. Single amino acid mutations of AVR2 in race 3 of the pathogen abolish recognition by I2 and overcome I2 mediated disease resistance (Houterman, Ma et al. 2009, Takken and Rep 2010). A current challenge is to breed *R* genes that respond to the stealthy AVR2 variants.

*S. demissum R3a* is one of the first late blight *R* genes to be bred in cultivated potato (Hawkes 1990, Gebhardt and Valkonen 2001, Huang, van der Vossen et al. 2005). The R3a immune receptor strongly responds to the *P. infestans* RXLR-type host-translocated AVR3a^K80/I103^ effector (referred to as AVR3a^KI^) but weakly to AVR3a^E80/M103^ (AVR3a^EM^), which differs in only two amino acids in the mature protein (Armstrong, Whisson et al. 2005, Bos, Kanneganti et al. 2006). *P. infestans* strains that carry *Avr3a*^*KI*^, either in a homozygous or heterozygous configuration, are avirulent on *R3a* potatoes (Armstrong, Whisson et al. 2005, Bos, Kanneganti et al. 2006, Bos, Chaparro-Garcia et al. 2009, Chapman, Stevens et al. 2014, Segretin, Pais et al. 2014). *P. infestans* strains homozygous for *Avr3a*^*EM*^ are virulent on *R3a* and have increased in frequency in *P. infestans* populations since the 19^th^ century, to become dominant in many potato growing regions of the world (Cooke, Cano et al. 2012, Chowdappa, Kumar et al. 2013, Li, van der Lee et al. 2013, Yoshida, Schuenemann et al. 2013, Yoshida, Burbano et al. 2014). This prompted Segretin et al. to expand the response spectrum of R3a to AVR3a^EM^ by random mutagenesis. Eight single-amino acid mutants, termed R3a+, gained response to AVR3a^EM^ while retaining the ability to respond to AVR3a^KI^ (Segretin, Pais et al. 2014). Six of these R3a+ mutations locate to the LRR domain, one in the NB-ARC and one in the CC. The two mutants in the CC and NB-ARC domains, R3a^I148F^ and R3a^N336Y^, respectively, showed further gain-of-response to PcAVR3a4, an AVR3a homolog from the pepper pathogen *Phytophthora capsici*. Segretin *et al*. proposed that these mutants are sensitized, “trigger-happy” mutants, with a lower threshold for activation of the NLR receptor, therefore conferring enhanced response to weak elicitors (Segretin, Pais et al. 2014). However, the R3a+ mutants, and similar multiple site R3a mutants described by Chapman et al. (Chapman, Stevens et al. 2014), did not translate in enhanced resistance to *P. infestans* isolates homozygous for the *Avr3a*^*EM*^ allele, indicating that the observed gain-of-response in the cell death assay was insufficient for late blight resistance.

In this study, we took advantage of the relatively high sequence similarity between *S. demissum* R3a and its tomato ortholog I2 to test the extent to which gain-of-function mutants identified in one NLR can be transferred to a homologous receptor from another plant species. This study was based on the observation that two residues mutated in the R3a+ mutants, R3a^I148F^ and R3a^N336Y^, are conserved in I2 (Huang, van der Vossen et al. 2005), and our hypothesis that these residues are hotspots for sensitized phenotypes in the R3a/I2 class of NLRs. However, transfer of the two R3a+ mutations to I2 resulted in a loss-of-response for I2^I141F^ and autoactivity for I2^N330Y^. We then reasoned that other amino acid substitutions at the same positions might yield the desired phenotype, and therefore generated I2 mutant proteins carrying all possible amino acids in positions 141 and 330. Remarkably, one mutant, I2^I141N^, displayed expanded response to both AVR3a^KI^ and AVR3a^EM^, and conferred partial resistance to *P. infestans* strains carrying either of the AVR3a variants. In addition, I2^I141N^ responded to two stealthy *F. oxysporum f. sp. lycopersici* AVR2 effector variants that evade response by the wild-type I2 receptor. This work demonstrates that synthetic NLR receptors can confer expanded pathogen effector and disease resistance spectra, and that knowledge on one NLR can be exploited to improve homologous NLRs from other plant species.

## Results

### I2 responds weakly to AVR3a^KI^

Given that *I2* and *R3a* are orthologous genes, we first determined whether I2 responds to the two AVR3a variants from *P. infestans*. We co-expressed I2 with each of the AVR3a isoforms using *Agrobacterium tumefaciens*-mediated transient transformation (agroinfiltration) in the model plant *Nicotiana benthamiana* and scored for hypersensitive cell death phenotypes. We found that I2 responds weakly to AVR3a^KI^ (HR index typically between 0.5 and 1.2 on a scale of 0 to 3 at 6 days post-infiltration (dpi)) (Fig. 1 and S1). Occasionally, I2 showed a weak response to AVR3a^EM^ (HR index less than 0.5) (Fig. 1). Both AVR3a variants were expressed alone in *N. benthamiana* to exclude any possibility of weak autoactivity of these constructs (Fig. S2). No cell death reaction was seen upon transient expression of either AVR3a variant (Fig. S2), suggesting that the I2/AVR3a cell death is specific. These findings, together with the high amino acid sequence similarity between R3a and I2, prompted us to test the possibility that transferring previously identified R3a+ mutations to I2 expands the response profile of this NLR receptor.

**Figure 1:**
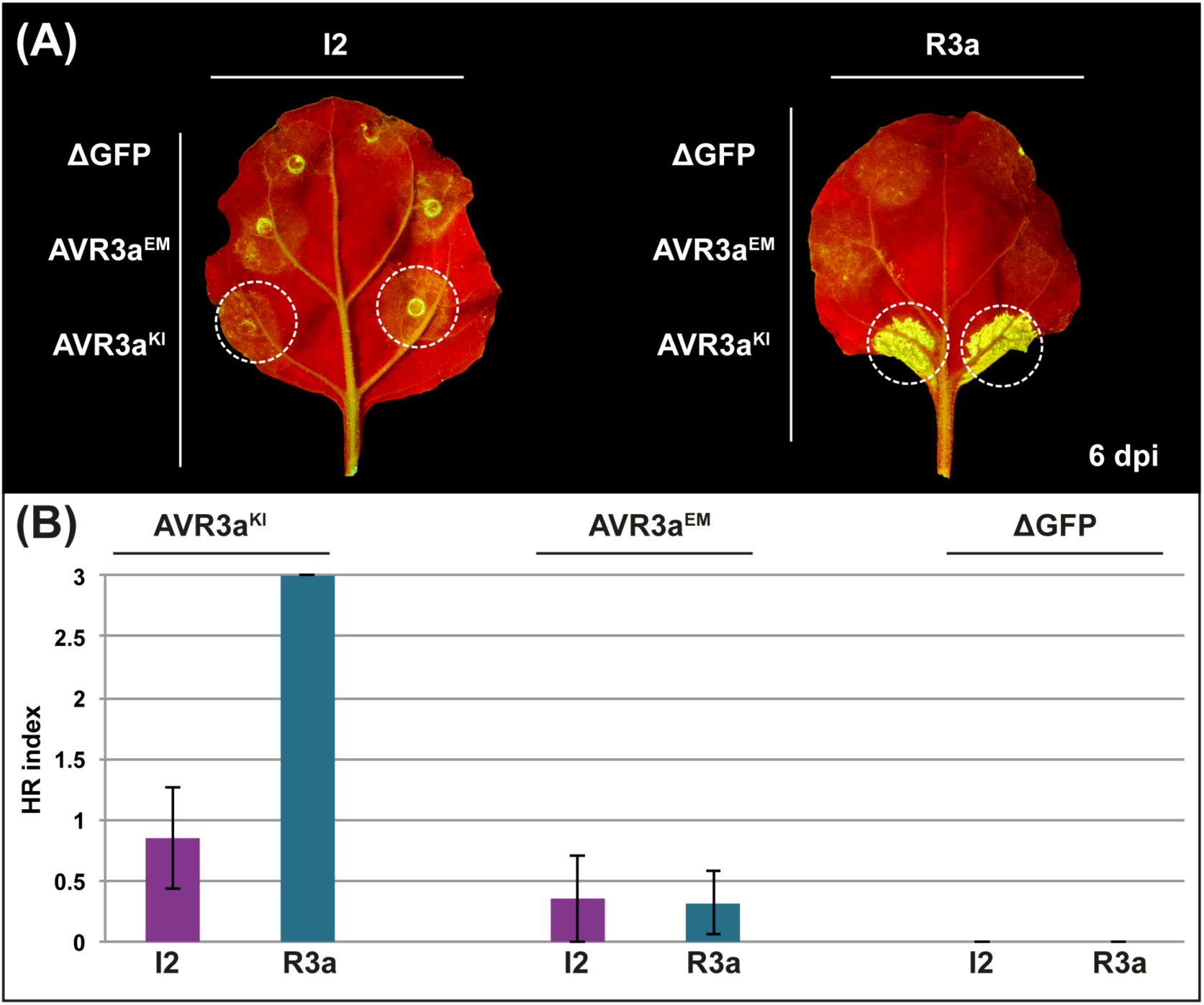
The tomato resistance protein I2 responds to AVR3a^KI^ from *Phytophthora infestans*. A, Hypersensitive response (HR) phenotypes of wild-type I2 after co-expression with the *P. infestans* AVR3a variants in *Nicotiana benthamiana* leaves. Wild-type *I2* was under transcriptional control of the *Cauliflower mosaic virus* 35S promoter and AVR3a^KI^, AVR3a^EM^ or a truncated version of green fluorescent protein (ΔGFP) were expressed from a *Potato Virus X* (PVX)-based vector. The picture was taken at 6 days post-infiltration (dpi). B, HR indices corresponding to the experiment described in A. Values scored at 6 dpi are plotted. Bars represent the average of 20 replicas for each combination of constructs; error bars represent standard deviation.

### Homology models of the NB-ARC domain highlights conserved R3a+ residues

Although the overall similarity between R3a and I2 is relatively high (83% amino acid similarity), the NB-ARC domain displays the highest sequence conservation (Huang, van der Vossen et al. 2005) (Fig. 2A). We generated structural models of the conserved CC and NB-ARC domains using four on-line prediction programs (IntFOLD2, iTASSER, Phyre2 and SWISS-MODEL).

**Figure 2:**
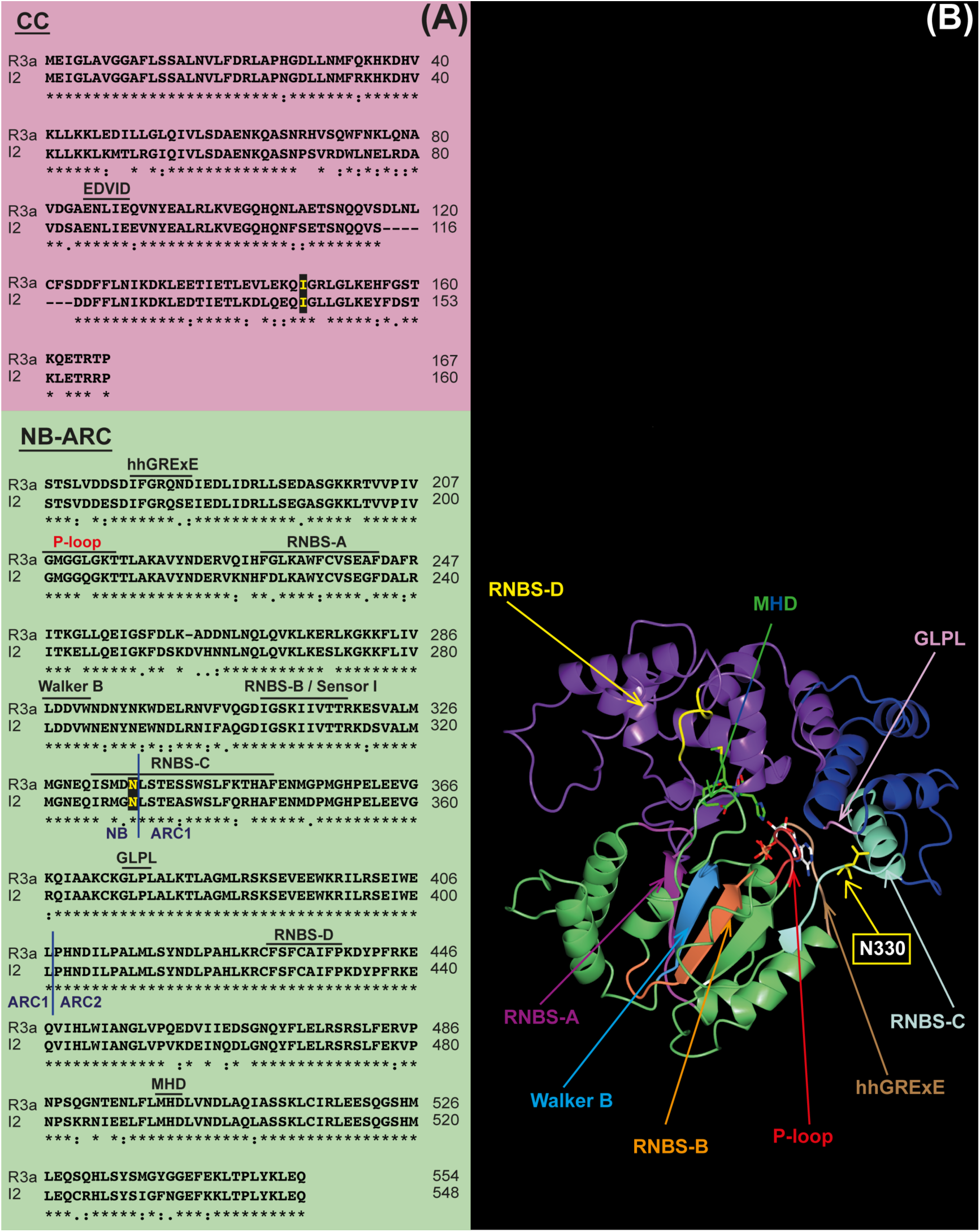
I2 and R3a are highly similar in the N-terminal region. A, Protein sequence alignment of the CC and NB-ARC domains of I2 and R3a. The amino acid positions that were subjected to mutation are highlighted in yellow. Conserved R-protein motifs in the NB-ARC domain are overlayed according to Ooijen *et al*, (Ooijen, Burg et al. 2007). Blue lines delimit the 3 subdomains of the NB-ARC: NB, ARC1 and ARC2. B, Structural model of the NB-ARC domain of I2. Models were generated by remote homology modeling using the INTFold2 server (Buenavista, Roche et al. 2012, McGuffin, Atkins et al. 2015). The NB-ARC is shown in three different colours depending on the subdomain: NB (green), ARC1 (purple), ARC2 (blue). Amino acids corresponding to the single-residue I2 mutations are shown in yellow stick representation. The carbon atoms of ATP are shown in white. The locations of NLR-protein motifs are indicated with arrows.

For the CC domain, three of the prediction programs (iTASSER, Phyre2 and SWISS-MODEL) identified the CC domain of the barley CNL protein MLA10 as the top-scoring template for modelling of the I2 CC-domain. Interestingly, IntFOLD2 (Buenavista, Roche et al. 2012, McGuffin, Atkins et al. 2015) produced a model based on the CC domain of the potato CNL protein Rx, which adopts a distinct structure, quite different at the three-dimensional level to MLA10. Despite these different models (either of which could be valid), in each of the structures the I2 I141 residue is consistently positioned to the C-terminus of the defined coiled-coil units and is therefore predicted to be located at an inter-domain region between the CC and NB-ARC domains (Fig. S3).

For the NB-ARC domain, the structures of the equivalent regions of Apaf-1 (Vaughn, Rodriguez et al. 1999), DARK (Pang, Bai et al. 2015) and CED-4 (Yan, Chai et al. 2005) were consistently used as templates by the programs for modelling, with high confidence scores. We choose to use the models based on Apaf-1, as the crystal structure of this template has been determined to the highest resolution and the residues forming the ATPase active site are appropriately positioned. As would be predicted from the modelling of the R3a NB-ARC domain (Segretin, Pais et al. 2014), the I2 N330 position maps to the junction of the NB and ARC1 regions (Fig 2B), near the NB binding site and adjacent regions that have been implicated in nucleotide exchange events in plant NB-LRR proteins and associated with activity (Van Ooijen, Mayr et al. 2008).

The structural models indicate that the two gain-of-function positions in the I2 CC and NB-ARC domain occur in inter-domain regions and mutations at these positions may affect the overall conformational dynamics of the protein. We therefore proceeded with the generation of I2 mutants in these positions.

### The I2^I141F^ and I2^N330Y^ mutants are nonfunctional and autoactive, respectively

We generated mutants of I2 that are equivalent to R3a^I148F^ and R3a^N336Y^, the R3a+ mutations in the CC and NB-ARC domains. We assayed these I2 mutants, I2^I141F^ and I2^N330Y^, with AVR3a^KI^ and AVR3a^EM^ using agroinfiltration in *N. benthamiana*. I2^I141F^ showed a loss-of-response as it did not trigger any cell death when co-expressed with either one of the AVR3a isoforms (Fig. 3). I2^N330Y^ was autoactive, it triggered strong cell death (HR index 3) even in the absence of effector proteins (Fig. 3). These results indicate that the specific R3a+ amino acid substitutions do not confer desirable phenotypes to I2, but highlight the importance of these two positions for I2 activity.

**Figure 3:**
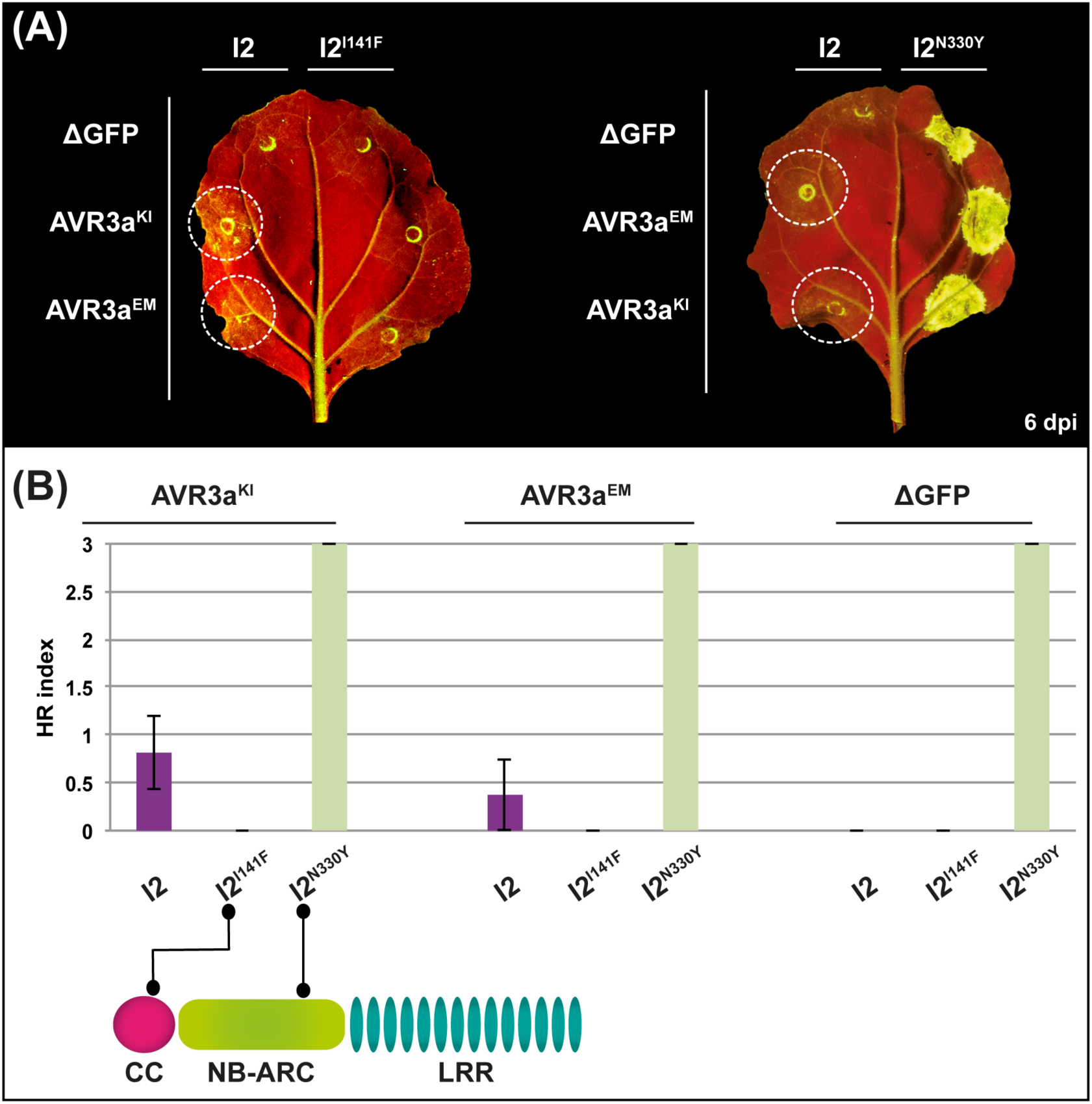
I2 mutants carrying the precise R3a+ singe-amino acid mutations in the CC and NB-ARC domains show a loss-of-response and an autoactive phenotype respectively. A, Hypersensitive response (HR) phenotypes of wild-type I2, I2^I141F^ and I2^N330Y^ after co-expression with the *Phytophthora infestans* AVR3a variants in *Nicotiana benthamiana* leaves. Wild-type *I2* and *I2* mutants were under transcriptional control of the *Cauliflower mosaic virus* 35S promoter and AVR3a^KI^, AVR3a^EM^ or a truncated version of green fluorescent protein (ΔGFP) were expressed from a *Potato Virus X* (PVX)-based vector. The pictures were taken at 6 days post-infiltration (dpi). B, HR indices corresponding to the experiment described in A. Values scored at 6 dpi are plotted. The cartoon indicates the approximate positions of the mutations. Bars represent the average of 16 replicas for each combination of constructs; error bars represent standard deviation.

### I2 mutants at positions 141 and 330

We hypothesized that additional amino acid substitutions at I2 positions 141 and 330 might yield an expanded response to AVR3a. To test this, we randomly mutagenized I2 codons 141 and 330 (Fig. 4). Next, we cloned mutagenized *I2* molecules in the binary vector pk7WG2 (Karimi, Inze et al. 2002) under the transcriptional control of the *Cauliflower mosaic virus* 35S promoter, and transformed them into *A. tumefaciens.* We obtained mutants representing all different amino acids for position 330, whereas we only generated a library with 12 amino acids for position 141. We screened the mutant clones by co-expression with the AVR3a isoforms in *N. benthamiana* and grouped the cell death phenotypes in four categories: strongly autoactive (HR index = 3 in the absence of effectors), loss-of-response (no HR phenotype), similar profile to wild-type I2, and gain-of-response (expanded response to AVR3a^EM^). Substitutions of the Asn in position 330 with Cys, His, Leu, Tyr, Arg, Thr, Ser, Val, Trp, Phe, Met and Ile resulted in autoactive mutants; Glu, Pro and Asp gave a loss-of-response phenotype and Lys, Gln, and Ala had a similar response profile to the wild-type I2 (Fig. 4). Substitution of the Ile in position 141 with Tyr resulted in an autoactive mutant; Arg, Pro, Val, Ala, Lys, Leu, Thr and Asp showed a loss-of-response phenotype. Interestingly, I2^I141N^, which carries a substitution of Ile to Asn in position 141, responded more strongly to AVR3a^EM^ compared to the wild-type I2 (Fig. 5).

**Figure 4:**
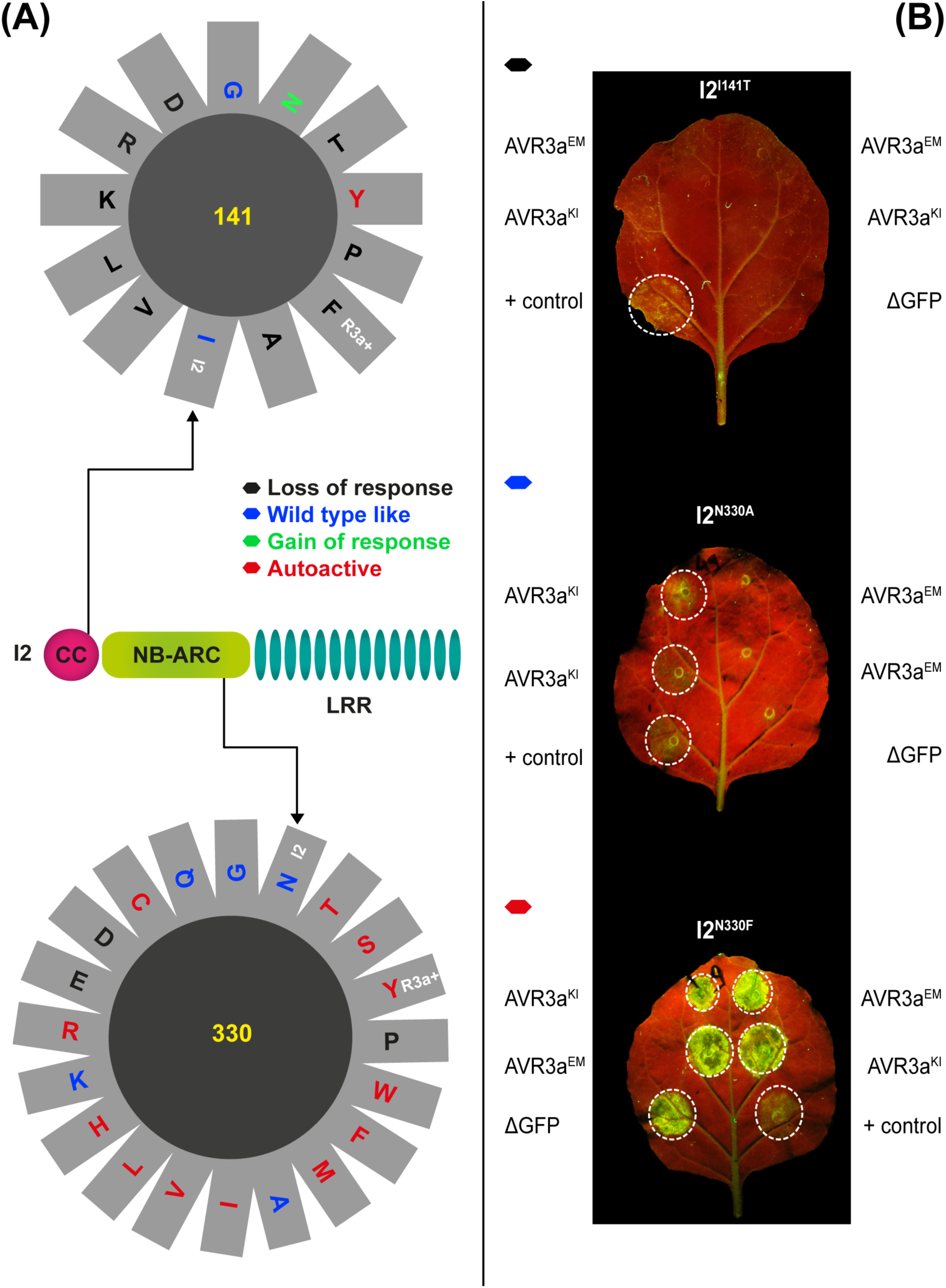
I2 mutant libraries carrying randomized amino acids at positions 141 and 330. The full length *I2* coding sequence (3,801 bp) was used as template for a polymerase chain reaction-based mutagenesis targeted to the corresponding coding sequence positions. Amplification products were cloned in the binary vector pK7WG2 (under transcriptional control of the *Cauliflower mosaic virus* 35S promoter) and transformed into *Agrobacterium tumefaciens*. The library was screened for gain-of-response phenotypes by co-agroinfiltration in *Nicotiana benthamiana* of the I2 mutant clones with *Phytophthora infestans* AVR3a^KI^ and AVR3a^EM^ expressed from a *Potato virus X* (PVX)-based vector. A, Amino acid substitutions are coloured according to the response phenotypes obtained: black (loss-of-response), blue (wild-type like), green (gain-of-response), red (autoactive). Wild-type amino acids at these positions and amino acids corresponding to the R3a+ mutants are indicated by the I2 and R3a+ labels respectively. B, Hypersensitive response (HR) phenotypes of selected I2 mutants generated in this library (I2^I141T^, I2^N330A^, I2^N330F^) after co-expression with the *Phytophthora infestans* AVR3a variants in *N. benthamiana* leaves, representing a loss-of-response, a wild-type like and an autoactive phenotypic example respectively. Positive control (+), represents I2 wild-type co-expressed with AVR3a^KI^ in each leaf.

**Figure 5:**
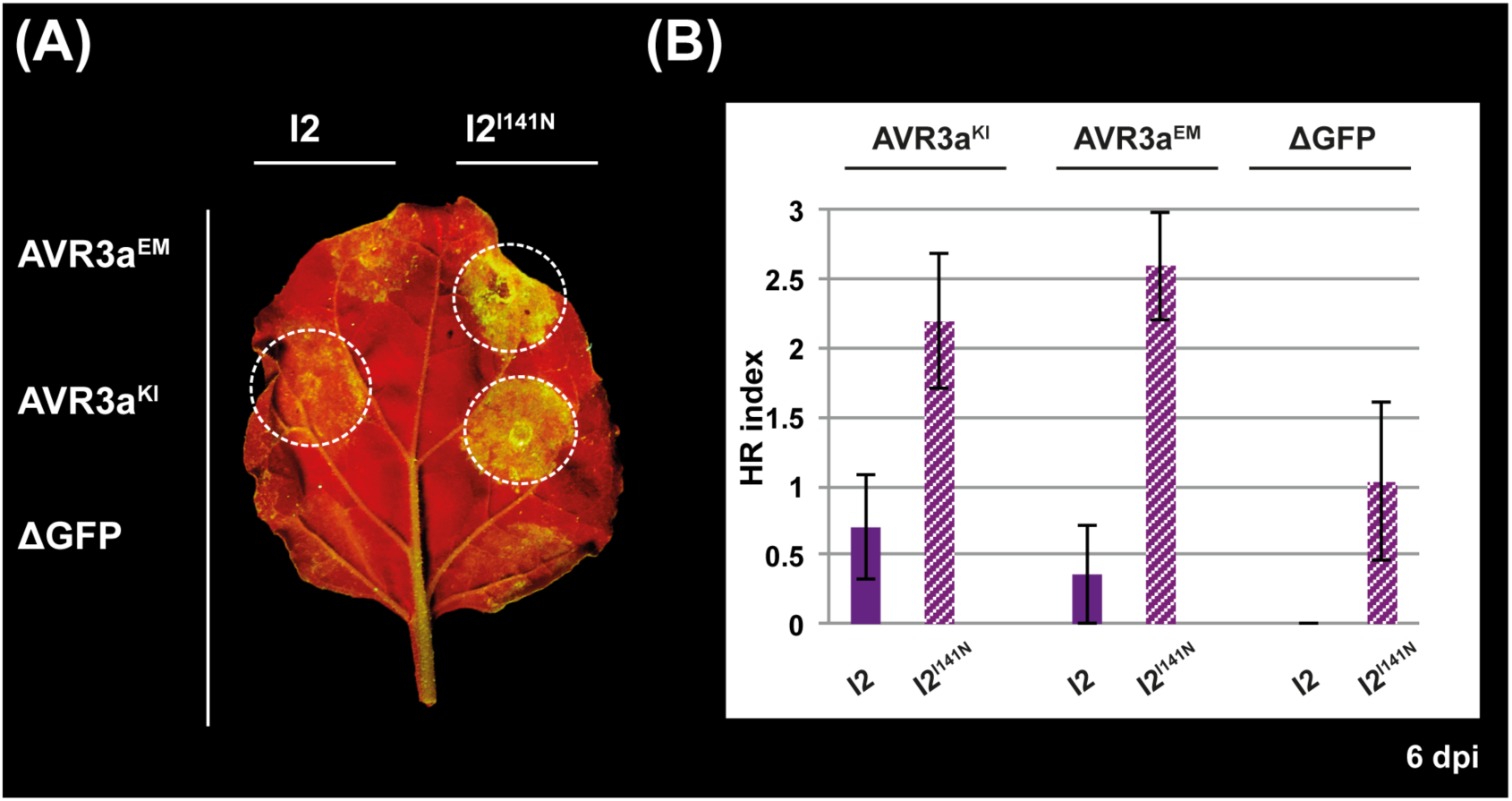
I2^I141N^ responds to AVR3a^EM^ while retaining the response to the AVR3a^KI^ isoform. A, Hypersensitive response (HR) phenotypes of wild-type I2 and I2^I141N^ after co-expression with the *Phytophthora infestans* AVR3a variants in *Nicotiana benthamiana* leaves. The wild-type and mutant *I2* were under transcriptional control of the *Cauliflower mosaic virus* 35S promoter and AVR3a^KI^, AVR3a^EM^ or a truncated version of green fluorescent protein (ΔGFP) were expressed from a *Potato Virus X* (PVX)-based vector. The picture was taken at 6 days post-infiltration (dpi). B, HR indices corresponding to the experiment described in A. Values scored at 6 dpi are plotted. Bars represent the average of 20 replicas for each combination of constructs; error bars represent standard deviation.

### A single amino acid change expands the response of I2 to AVR3a^EM^

To determine the extent to which the single-site I141N mutation in the CC domain of I2 expands its response profile to the AVR3a^EM^ variant, we co-expressed I2^I141N^ with both AVR3a isoforms using *N. benthamiana* agroinfiltration. I2^I141N^ responded to AVR3a^EM^ (HR index between 2.5 and 3 in a scale of 0 to 3), and also exhibited a stronger response to AVR3a^KI^ when compared to wild-type I2 (HR index ~2 vs 0.7 for the wild-type) (Fig. 5). I2^I141N^ also exhibited a weak response in the absence of effector (HR index ~1), which is lower than the response to the two AVR3a variants. These results indicate that a single-amino acid change in the CC domain of I2 (I141N) expanded its response profile to AVR3a^EM^ and increased response of the receptor to the AVR3a^KI^ isoform.

To examine whether the activity of the I2^I141N^ mutant protein extends to additional AVR3a variants besides AVR3a^KI^ and AVR3a^EM^, we tested both I2 wild-type and I2^I141N^ mutant for their response to selected AVR3a mutants with amino acid substitutions at position 80 in AVR3a^K80X/I103^ and AVR3a^E80X/M103^ (Bos, Chaparro-Garcia et al. 2009). I2^I141N^ showed expanded response to all of the mutants tested (Fig. S4).

### I2^I141N^ protein accumulates at different levels compared to wild-type I2

To determine the effect of the I141N mutation on the stability of the I2 protein, we performed western blot assays using a polyclonal antibody raised against the CC domain of R3a. We collected samples at 2, 3 and 5 dpi, as we reasoned that this range of time points would allow us to detect potential differences in the protein levels between I2 and I2^I141N^. Interestingly, at 2 and 5 dpi, wild-type I2 yielded a more intense signal than the I2^I141N^ mutant (Fig. 6). The loss-of-response mutant protein I2^I141T^ was used as an additional control in this experiment and only yielded a weak signal at 5 dpi (Fig. 6). Additional analysis on the remaining loss-of-response mutants at the N-terminal part of I2, showed that all of them are expressed (Fig. S5). Of the I2 loss-of-response mutants tested I2^I141D^, I2^I141T^, I2^I141L^, I2^I141K^, I2^I141V^, I2^I141F^ and I2^N330E^ all accumulated to lower level than the wild-type I2 protein (Fig. S5).

**Figure 6:**
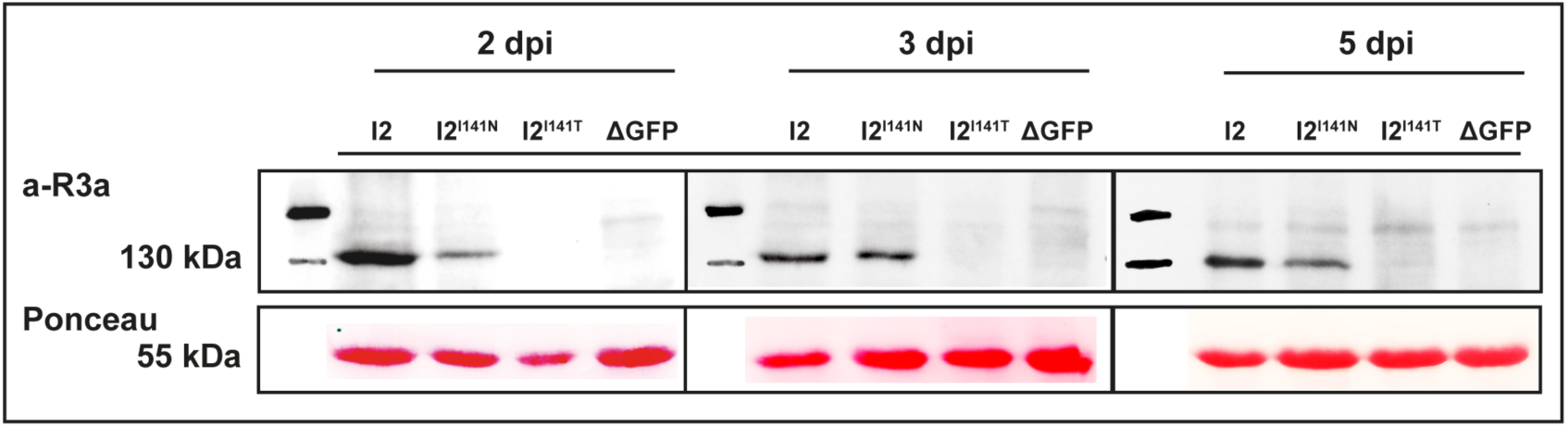
The I2^I141N^ gain-of-response mutant has an altered stability compared to wild-type I2 *in planta*. *Nicotiana benthamiana* leaves were agroinfiltrated with constructs to express wild-type and mutant I2 proteins or a truncated version of green fluorescent protein (ΔGFP). Total protein extracts from leaves sampled at 2, 3 and 5 days post-infiltration (dpi) were subjected to SDS-PAGE followed by Immunoblotting with a polyclonal antibody raised against the CC domain of R3a (a-R3a, upper panel). A band of approximately 145 kDa corresponding to I2 was present in total extracts of the leaves infiltrated with the *I2* and *I2*^*I141N*^ constructs. Ponceau S staining of Rubisco (lower panel) is shown as control of the amount of protein loaded and transferred in each lane. Sizes (in kDa) are indicated on the left. I2^I141T^ is a loss-of-response I2 mutant and was included as an additional negative control in this experiment. The different boxes indicate different blots.

### R3a^I148N^ has a response profile similar to that of wild-type R3a

To investigate whether the I141N mutation in the CC domain of I2 could also alter the response profile of R3a, we introduced this mutation in R3a to generate the R3a^I148N^ mutant. We then co–expressed R3a^I148N^ with the AVR3a variants using agroinfiltration assays in *N. benthamiana* and scored for the HR. The response of R3a^I148N^ to the two AVR3a isoforms was similar to that of wild-type R3a, as a potent hypersensitive response was detected only when the protein was co-expressed with AVR3a^KI^ (HR index ~3) (Fig. S6). As previously reported (Bos, Chaparro-Garcia et al. 2009) a weak response was sometimes observed when the wild-type R3a protein was co-expressed with the AVR3a^EM^ variant, a trend that was also observed for the R3a^I148N^ mutant (HR index < 0.2, Fig. S6). These results further indicate that, even though the 141/148 amino acid position in the CC domain is important for the activities of I2 and R3a, different amino acid substitutions are necessary to expand the response spectrum of these immune receptors.

### I2^I141N^ confers partial resistance to a *P. infestans* strain expressing AVR3a^EM^

To determine the degree to which the expanded response profile of I2^I141N^ translates into a wider resistance spectrum, we performed infection assays in *N. benthamiana* leaves expressing I2 and I2^I141N^ with different strains of *P. infestans*. We expressed I2 and I2^I141N^ using *A. tumefaciens* mediated transformation in leaves of three to four-week-old *N. benthamiana* plants. Approximately 15 hours after agroinfiltration, we inoculated infiltrated leaves with *P. infestans* zoospore suspensions. We first examined the effect of the wild-type I2 protein on *P. infestans* strain NL00228 homozygous for *Avr3a*^*KI*^ (Zhu, Li et al. 2012), or strain 88069 homozygous for *Avr3a*^*EM*^ (van West, de Jong et al. 1998). I2 expression restricted infection by *P. infestans* NL00228 as illustrated by smaller lesion sizes compared to those observed in leaves expressing a control construct (Fig. 7). In contrast, I2 did not impact infection by strain 88069 resulting in no significant differences in lesion sizes (Fig. 7). Remarkably, the I2^I141N^ mutant protein restricted infection by both strains of *P. infestans*. In more detail, the effect of I2^I141N^ on the *P. infestans* NL00228 strain was to a similar level to the wild-type I2 (Fig. 7). Restriction of *P. infestans* 88069 growth in leaves expressing I2^I141N^ was evident as smaller lesions developed compared to those developed in leaves expressing a control construct (Fig. 7 and S7). These findings suggest that the I2^I141N^ mutant has an expanded resistance profile to *P. infestans* relative to the I2 wild-type receptor.

**Figure 7:**
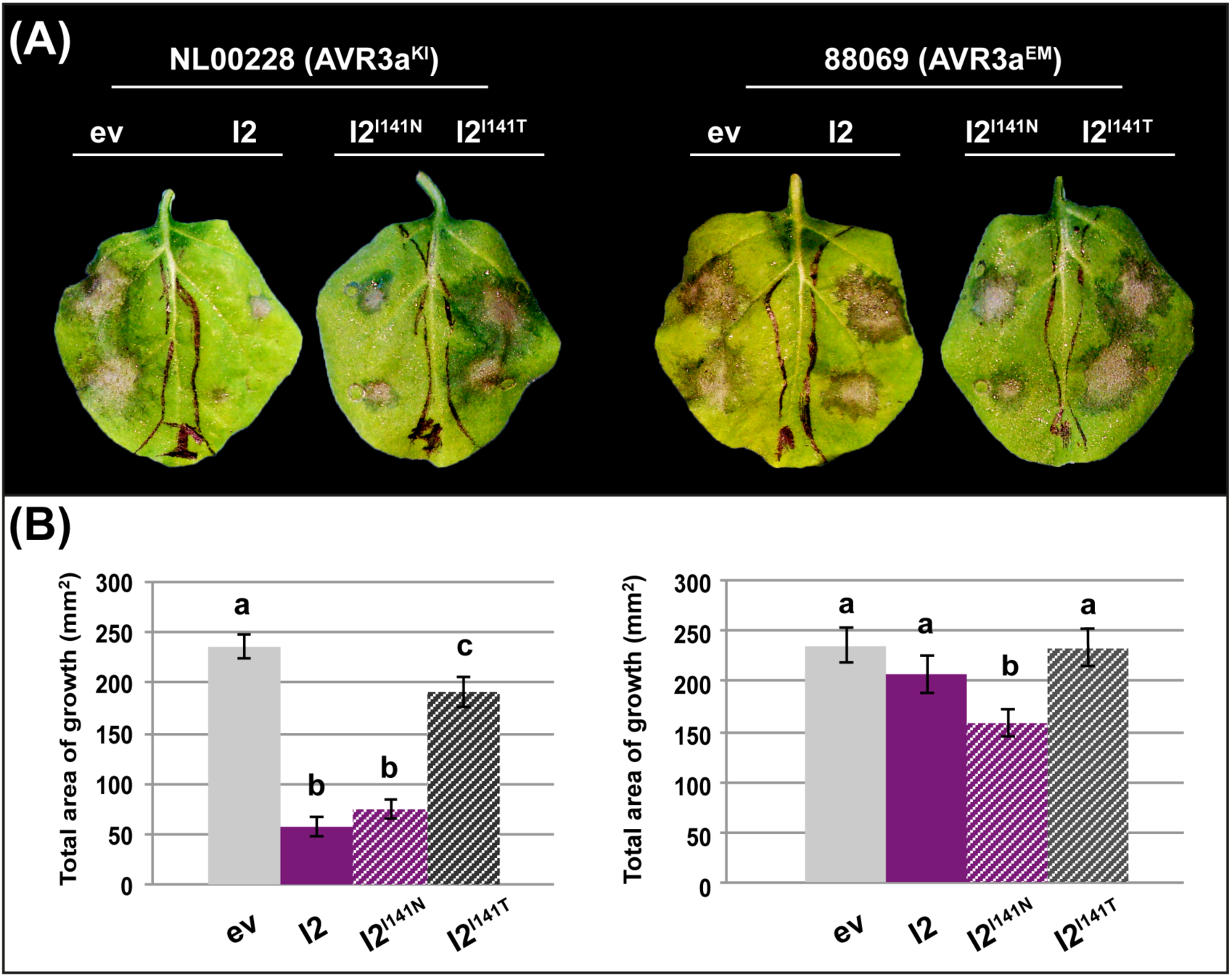
Wild-type I2 and I2^I141N^ confer partial resistance to strains of *Phytophthora infestans*. A, Wild-type I2 and I2 mutants were expressed in *Nicotiana benthamiana* leaves under transcriptional control of the *Cauliflower mosaic virus* 35S promoter. After approximately 15 hours, the infiltrated leaves were drop-inoculated with *P. infestans-*zoospore suspensions corresponding to the strains NL00228 (carrying AVR3a^KI^) and 88069 (carrying AVR3a^EM^). The pictures were taken at 6 days post-inoculation. B, The total area of *P. infestans* growth was determined with Image J software (Schneider, Rasband et al. 2012). Values corresponding to 6 days post-inoculation are plotted. Bars represent the average of 24 replicas for each treatment; error bars represent standard deviation. The experiment was performed 4 times and a representative repeat is shown. The Gateway binary vector pK7WG2 was included as negative control (empty vector, ev). I2^I141T^ is a loss-of-response I2 mutant and was used as an additional negative control in this experiment. Significant differences between the groups are indicated by letters and were determined in an analysis of variance (ANOVA) (p≤0.05).

### I2^I141N^ responds to *F. oxysporum f. sp. lycopersici* stealthy effectors AVR2^V>M^ and AVR2^R>H^

We also tested the extent to which I2^I141N^ has altered response to AVR2 effectors of *F. oxysporum f. sp. lycopersici* (Takken and Rep 2010). To achieve this, we co-expressed both I2 wild-type and I2^I141N^ mutant proteins with the AVR2 effector variants from *F. oxysporum f. sp. lycopersici* using agroinfiltration in *N. benthamiana*. I2^I141N^ responded to two out of the three stealthy AVR2 variants: AVR2^V>M^ (HR index ~3) and AVR2^R>H^ (HR index ~1.2), while retaining the capacity to respond to AVR2 from *F. oxysporum f. sp. lycopersici* race 2 (HR index ~2.5) (Fig. 8). These results indicate that the gain of response mutation identified based on one pathogen effector displays also increased response spectrum to effectors from another pathogen. The I2^I141N^ mutant of I2 has, therefore, expanded response to effectors from oomycetes and fungi, two phylogenetically unrelated plant pathogens.

**Figure 8:**
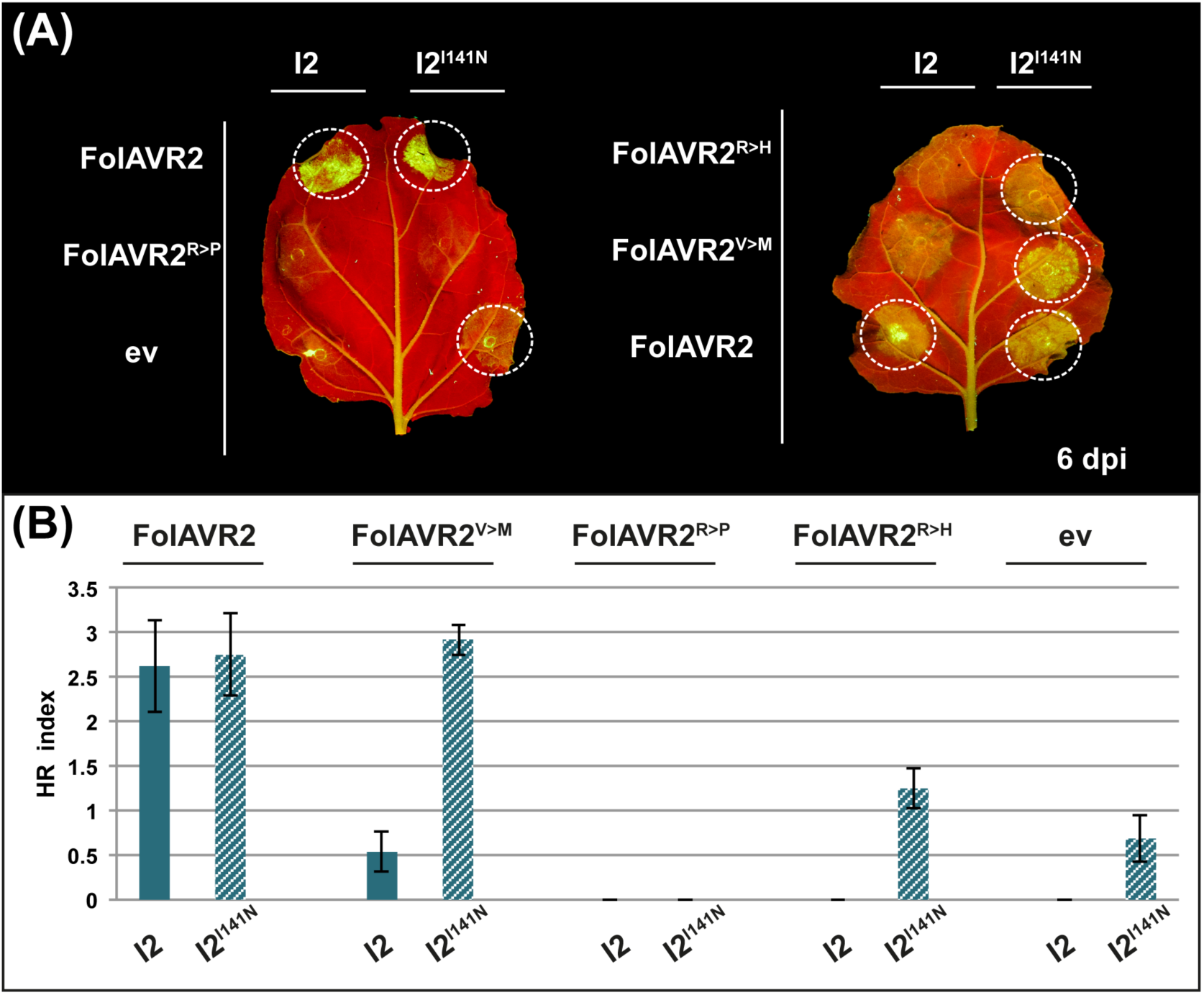
I2^I141N^ shows expanded response to *Fusarium oxysporum f. sp. lycopersici* (Fol) AVR2 variants from race 3. A, Hypersensitive response (HR) phenotypes of wild-type I2 and I2^I141N^ after co-expression with the FolAVR2 variants from race 3 in *Nicotiana benthamiana* leaves. Wild-type and mutant *I2* were under transcriptional control of the *Cauliflower mosaic virus* 35S promoter while FolAVR2, FolAVR2^V>M^, FolAVR2^R>P^ and FolAVR2^R>H^ were expressed from the binary vector CTAPi. The CTAPi empty vector (ev) was included as negative control. The picture was taken at 6 days post-infiltration (dpi). B, HR indices corresponding to the experiment described in A. Values scored at 6 dpi are plotted. Bars represent the average of 18 replicas for each combination of constructs; error bars represent standard deviation.

## Discussion

Tomato I2 and its potato ortholog R3a are two NLR immune receptors that have been previously described to mediate resistance to fungi and oomycetes respectively (Ori, Eshed et al. 1997, Simons, Groenendijk et al. 1998, Armstrong, Whisson et al. 2005, Bos, Kanneganti et al. 2006, Takken and Rep 2010). We discovered that I2 weakly responds to the AVR3a^KI^ effector from *P. infestans* and also confers resistance to a strain of this pathogen that expresses this effector (Fig. 1, Fig. 7). By using targeted mutagenesis, we recovered a mutant in the I2 protein, I2^I141N^, with enhanced response to both AVR3a^KI^ and the stealthy AVR3a^EM^ isoform that is produced by R3a-virulent races of *P. infestans* (Fig 5). Remarkably, I2^I141N^ conferred partial resistance to an R3a-breaking strain of *P. infestans* that expresses AVR3a^EM^ (Fig. 7). In addition, I2^I141N^ showed enhanced response to two out of three *F. oxysporum f. sp. lycopersici* AVR2 stealthy variants that evade response by the wild-type I2 (Fig. 8). These findings open up exciting perspectives for the budding field of synthetic NLR-mediated disease resistance (Farnham and Baulcombe 2006, Harris, Slootweg et al. 2013, Segretin, Pais et al. 2014). First, knowledge generated on one NLR receptor can be exploited to improve homologous NLRs from other plant species. Second, synthetic NLR immune receptors can be engineered to confer resistance to phylogenetically divergent pathogens. Overall, these approaches can be applied to develop synthetic immune receptors that are beneficial to agriculture.

In this study we focused on two previously identified mutations at the N terminal part of R3a (R3a^I148F^ and R3a^N336Y^) and transferred them to I2 based on the high level of sequence similarity the two proteins have at this region. The I2 mutants generated resulted in a loss-of-response (I2^I141F^) and autoactive phenotype (I2^N330Y^), respectively. Our knowledge of the underlying mechanisms in the I2 and R3a systems is limited. However, we propose that previous screens (Chapman, Stevens et al. 2014, Segretin, Pais et al. 2014) have revealed amino acid positions that are important for the activity of these NLR receptors. One possibility is that the amino acids surrounding these two positions in the CC and NB-ARC domain of I2 (141 and 330), combined with the introduced substitutions, affect protein activity by preventing interaction with downstream signalling components (I2^I141F^) or by changing the overall conformation of the protein leading to a constitutive activated state (I2^N330Y^). Based on our structural modelling, the position 141 of I2 is located after the structurally defined CC domain, at an inter-domain region between the CC and NB-ARC regions (Fig. S3). Therefore, it is possible that a change in this residue perturbs domain-domain interactions, affecting the function of this region and interfering with the overall performance of the protein. Previous studies have highlighted the involvement of the CC domain in pathogen perception and downstream signaling (Rairdan and Moffett 2006, Maekawa, Cheng et al. 2011, Chen, Liu et al. 2012, Hao, Collier et al. 2013). Rairdan *et al*., have previously proposed that motifs in the CC domain, including the highly conserved EDVID, mediate interactions important for the function of the protein (Rairdan, Collier et al. 2008). However, position 141 in the I2 CC domain doesn’t appear to be close to the EDVID motif (Fig. 2A), therefore we cannot be sure about the mechanism underlying the mutation in that position. In the same study, Rairdan *et al*., further suggest a model where the signaling activity of the NB domain is defined by the joint function of both the CC and LRR domains and that this regulation is recognition dependent (Rairdan, Collier et al. 2008). Therefore, it is also possible that the specific amino acid change in the CC domain perturbs intramolecular interactions with the LRR domain thus enhancing the signaling activity of I2.

The position 330 is located near the predicted nucleotide-binding pocket of I2 (Fig. 2B). This region is thought to act as a switch between the active and inactive state of NLR proteins (Tameling, Vossen et al. 2006). In a previous study, Harris et al. showed that single-amino acid mutations near the nucleotide-binding pocket of the potato NLR protein Rx enhanced the expanded response conferred by a single mutation in the LRR domain, resulting in resistance to PopMV (Harris, Slootweg et al. 2013). However, saturation mutagenesis in position 330 in I2 did not result in expanded response phenotypes. It is possible that mutations in this position affect the affinity or catalysis of the ATP/ADP nucleotide. Further analysis is required before any firm conclusions can be drawn.

Stirnweis *et al*., identified two positions in the NB-ARC domain of the wheat powdery mildew resistance gene *Pm3*, that when mutated enhance the ability of the protein to trigger cell death in *N. benthamiana* and also result in expanded resistance of the *Pm3f* allele in wheat. They further showed that the same mutations affected the activity of another CNL rice immune receptor, Bph14, highlighting the importance of these two amino acids in a distantly related protein (Stirnweis, Milani et al. 2014). In addition, Ashikawa *et al*, working on two members of the Pik locus in rice, the broad-spectrum *Pikm1-TS* and narrow spectrum *Pik1-KA* gene, generated chimeras to identify which domains define the recognition specificity. They found that a combination of the CC-NB of Pikm1-TS and the LRR of Pik1-KA was sufficient for resistance to a blast fungus isolate virulent only to Pik1-KA. Their results suggested that one or more of the polymorphic amino acids in the CC and/or NBS domain are responsible for the recognition specificity (Ashikawa 2012). Overall, these studies support the view that the N-terminal part of NLR protein is implicated in the perception of the pathogen. In conclusion, although our I2^I141N^ mutant appears to be sensitized it may also have altered specificity. Future work aiming at identifying the mechanisms mediating the observed expanded response of the I2^I141N^ mutant protein and also addressing whether this response translates to resistance are required.

Even though transfer to I2 of the precise mutations previously identified in R3a did not yield the desirable phenotype, they helped us focus our screens for improved I2 mutants. The I2^I141N^ mutant identified in this study has expanded response to both AVR3a isoforms from *P. infestans* (Fig. 5) and also to two AVR2 variants from *F. oxysporum f. sp. lycopersici* race 3 (Fig. 8). However, this mutant also shows a weak HR in the absence of effectors, which suggests that it has increased activation sensitivity. Most likely, this sensitized “trigger-happy” mutant has a lower threshold for activation compared to wild-type I2 and is more easily activated by weak elicitors. The altered stability of I2^I141N^ (Fig. 6) is another indication that this mutation affects the overall conformation of the protein that could make it easier for the protein to get activated (Schornack 2006). Another possibility is that the mutated protein has an altered ATP binding ability. In the wild-type I2 protein, both forms (ATP-bound, “on”; ADP-bound, “off”) are kept in a dynamic equilibrium and in the absence of an elicitor I2 would predominantly be inactive. In a previous study, an autoactive version of I2 was shown to be impaired in nucleotide hydrolysis (Tameling, Vossen et al. 2006). A similar mechanism was proposed for an autoactive version of the flax M NLR protein, which shows increased preference for binding ATP (Williams, Sornaraj et al. 2011). Further biochemical tests on I2^I141N^ are required to assess whether this mutation in the N-terminal part of I2 alters the ability of the protein to bind ATP, modifying the threshold for activation.

The pathogenicity assays showed that I2 confers resistance to a *P. infestans* isolate homozygous for the *Avr3a*^*KI*^ allele (Fig. 7) suggesting that tomato lines carrying this gene may confer strain-specific resistance to *P. infestans*. However, the level of HR observed between I2 and AVR3a^KI^ was relatively weak therefore we cannot rule out that other effectors present in this *P. infestans* determine the observed I2-mediate resistance. Most late blight lineages infect both potato and tomato (Goodwin, Smart et al. 1998), although some lineages primarily infect tomato and are not likely to cause disease on potato plants in the field (Hu, Perez et al. 2012). In a later study, Armstrong et al, screened a selection of 45 *P. infestans* isolates from Canada, Mexico, US, Argentina, Ecuador, Denmark, Netherlands and Scotland and found that 40 of them were homozygous for the *Avr3a*^*EM*^ allele, 14 carried both *AVR3a*^*KI*^ and *AVR3a*^*EM*^ while only 1 was homozygous for *Avr3a*^*KI*^ (Armstrong, Whisson et al. 2005). Yoshida et al, after comparing the genomes of 11 ancient *P. infestans* strains with 15 modern isolates, revealed that the *AVR3a*^*KI*^ allele was dominant in ancient populations whereas both AVR3a alleles are found in modern isolates, with lineages homozygous for *Avr3a*^*EM*^ being more common (Yoshida, Schuenemann et al. 2013). Even though in many *P. infestans* populations the *AVR3a*^*EM*^ allele dominates, epidemics are often caused by clonal lineages. *I2*-expressing tomatoes may provide some degree of resistance to epidemics caused by strains homozygous for *Avr3a*^*KI*^ however these races are rare in modern *P. infestans* populations. Therefore, improved mutants with a broader spectrum of resistance to the dominant *AVR3a*^*EM*^ carrying *P. infestans* races are needed. The R3a+ mutants previously identified failed to provide resistance against *P. infestans* (Chapman, Stevens et al. 2014, Segretin, Pais et al. 2014). Remarkably, the I2^I141N^ mutant identified in this study gives partial resistance to *P. infestans* strains carrying either of the *AVR3a* variants in the transient assays used (Fig. 7). These results suggest that tomato lines carrying the I2^I141N^ mutant could confer resistance against *P. infestans*.

*NLR* genes constitute a useful tool for generating sustainable disease resistant crops. Traditional breeding approaches based on these genes have been used extensively in tomato (Andolfo, Jupe et al. 2014). Recent transgenic strategies also allow the efficient transfer of *R* genes between plant species (Wulff, Horvath et al. 2011, Horvath, Stall et al. 2012, Narusaka, Kubo et al. 2013). However, the deployment of *NLR* genes through both conventional breeding and transgenic approaches is hampered by the low occurrence of *R* genes with the useful response specificities. In this study, we show that a synthetically generated NLR mutant protein has expanded response specificities towards two different pathogens. This approach could prove useful for designing and engineering NLRs with novel activities, where mutants identified in one gene could be transferred to homologs. Similar to *I2* and *R3a* which share a high sequence homology, other *R* genes with high sequence similarity could benefit from this approach. Such examples are *RPP8*, *RCY1* and *HRT* from *Arabidopsis thaliana*, *Rx* and *Gpa2* from potato, *Mi-1.2* and *Rpi-blb2* from tomato and *Solanum bulbocastanum*, respectively. Random mutagenesis screens like the one leading to the identification of the R3a+ mutants (Segretin, Pais et al. 2014) have been extremely useful to determine critical positions affecting the response profile of *R* proteins. Recently, the advent of genome editing in plants (Feng, Zhang et al. 2013, Jiang, Zhou et al. 2013, Li, Norville et al. 2013, Mao, Zhang et al. 2013, Miao, Guo et al. 2013, Nekrasov, Staskawicz et al. 2013, Shan, Wang et al. 2013, Xie and Yang 2013) has opened a completely new era in the field of plant biotechnology. The application of genome editing to generate synthetic *R* genes with expanded response specificities could significantly broaden the perspectives for breeding crop plants with a far-ranging resistance.

## Materials and Methods

### Microbial strains and growth conditions

*A. tumefaciens* strain GV3101 (Hellens, Mullineaux et al. 2000) and *E. coli* TOP10 (Invitrogen) were used in molecular cloning experiments and library construction. These strains were cultured as previously described (Sambrook and Russell 2001), at 28°C (for GV3101) or 37°C (for TOP10) in Luria-Bertani (LB) media supplemented with the appropriate antibiotics. DNA was electoporated into electrocompetent *A. tumefaciens* GV3101 or transformed by heat shock into competent *E. coli* TOP10 cells following standard protocols (Sambrook and Russell 2001). The following isolates were used in the *P. infestans* infection assays: NL00228 (Zhu, Li et al. 2012), 88069 (van West, de Jong et al. 1998) and 88069td (transgenic strain expressing the red fluorescent marker tandem-dimer RFP, known as tdTomato) (Giannakopoulou, Schornack et al. 2014). *Phytophthora* strains were grown on rye sucrose agar as previously described (Kamoun, van West et al. 1998), at 18°C in the dark.

### Targeted mutagenesis of *I2* and *R3a*

The *I2* (GenBank number KR108299) clone was used in this study. The *I2* open reading frame was amplified using primers I2G_F and I2G_R (Table S1) designed to add a 4-bp (CACC) sequence to the 5’ end of the *I2* sequence which is required for directional cloning using the pENTR/D-TOPO^®^ cloning kit (Invitrogen). I2 clones with single-residue mutations (I2^I141F^ and I2^N330Y^) were obtained by inverse PCR upon the pENTR::*I2* backbone with sets of primers (I141F_F, I141F_R and N330Y_F, N330Y_R) designed to introduce the desired mutations (Table S1). Random mutagenesis at positions 141 and 330 in the CC and NB-ARC domains of I2 respectively, was performed by inverse PCR upon the pENTR::*I2* backbone with sets of degenerate primers (I141X_F, I141F_R and N330X_F, N330Y_R) designed to introduce all possible sequence combinations at these positions (Table S1). The amplification products were cloned with the pENTR^(tm)^/D-TOPO® cloning kit following standard protocols (Xu and Li 2008). The Gateway-compatible binary vector pK7WG2 (Karimi, Inze et al. 2002) was used as destination for the mutagenized *I2* sequences. The R3a^I148N^ clone with a single-residue mutation (I148N) was obtained by GenScript Site-Directed Mutagenesis Service (Piscataway, NJ, U.S.A.) as described previously (Segretin, Pais et al. 2014). The binary vector pCBNptII-vnt1.1P/T-*R3a* (Segretin, Pais et al. 2014) was used as template and destination for the mutagenized *R3a* sequence.

### Sequence analyses

Plasmid DNA from the *I2* mutant clones was isolated and *I2* inserts were sequenced by GATC Sequencing Service (Constance, Germany) using several primers to allow full coverage. Base-calling and quality values were obtained using the Phred algorithm (Ewing, Hillier et al. 1998). Sequences were analyzed with MacVector 12.6 (Olson 1994).

### Agroinfiltration assays

Wild-type and mutant I2 and R3a clones were co-agroinfiltrated in *N. benthamiana* leaves to compare their relative response to different effector proteins. Each combination of wild-type or mutant R protein and effector protein (or negative control) was infiltrated as 16 to 20 replicates per experiment and every experiment was repeated at least 3 times. Briefly, 10 ml-LB media cultures with antibiotics [rifampicin at 50 mg/liter, gentamicin at 20 mg/liter, and spectinomycin at 50 mg/liter (I2 and AVR2 constructs) or ampicillin at 100 mg/liter (R3a and AVR3a constructs)] were inoculated with the library clones and grown at 28°C for 48 h (to reach an optical density at 600 nm [OD600] of 1 to 1.2). Cultures were pelleted by centrifugation (5 min at 3,500 rpm and 15°C) and resuspended with infiltration buffer (0.5% MS salts, 10 mM MES, 2% sucrose, and 200 μM acetosyringone, pH 5.6) to a final OD600 of 0.1. In all the agroinfiltration experiments pGR106-FLAG-AVR3a^KI^, pGR106-FLAG-AVR3a^EM^ (amino acids 23 to 147) constructs and pGR106-FLAG-AVR3a^E80X/M103^ or pGR106-FLAG-AVR3a^K80X/I103^ were used to express AVR3a mature proteins (without signal peptide) (Armstrong, Whisson et al. 2005, Bos, Kanneganti et al. 2006, Bos 2007, Bos, Chaparro-Garcia et al. 2009). *A. tumefaciens* GV3101 transformed with pGR106-ΔGFP [containing a truncated version of GFP as described by Bos and colleagues (Bos, Kanneganti et al. 2006)] was grown as the AVR3a clones. In addition, CTAPi-AVR2, CTAPi-AVR2^V>M^, CTAPi-AVR2^R>P^, CTAPi-AVR2^R>H^ (Houterman, Ma et al. 2009) were used to express AVR2 mature proteins (without signal peptide). *A. tumefaciens* GV3101 transformed with CTAPi (Rohila, Chen et al. 2004) was grown as the AVR2 clones. For transient co-expression of *R* gene clones and effector clones, the cells resuspended in infiltration buffer were mixed to have a final OD600 of 0.1 (R clones) and 0.5 (effector clones). Agroinfiltration experiments were performed on 4-week-old *N. benthamiana* plants. Plants were grown and maintained throughout the experiments in a controlled environment room with a temperature of 22 to 25°C and high light intensity. HR phenotype development was monitored from 3 to 7 dpi according to an arbitrary scale from 0 (no HR phenotype observed) to 3 (confluent necrosis on the infiltrated area) (Fig. S1). All the agroinfiltration assays were performed in Norwich, UK, apart from the assay shown in figure S4 that was performed in Argentina, US.

### Monitoring of HR development

*N. benthamiana* agroinfiltrated leaves were examined for HR-associated autofluorescence. For the assays performed in Norwich, we used a Nikon D4 camera with a 60mm macro lens (ISO set to 1250 or 1600) equipped with a yellow filter (Kodak Wratten No. 8). UV Blak-Ray® longwave (365nm) lamps B-100AP were spotlights and were moved around the subject during the exposure to give a more even illumination. For the assay described in figure S4 the following lamp, UVP.LLC, 8W model UVLMS-38, longwave (365nm) (Ultraviolet Products) was used and the camera contained no yellow filter. The autofluorescence under exposure to UV light, is associated with accumulation of phenolic compounds (Klement, Rudolph et al. 1990).

### Infection assays

*P. infestans* strains were grown on rye sucrose agar as described in (Kamoun, van West et al. 1998), at 18°C in the dark. Spores were harvested as previously described (Kamoun, van West et al. 1998, Schornack, van Damme et al. 2010) and diluted to 50,000 spores/ml. Droplets of 10 μl were placed onto the abaxial side of 4-week-old detached *N. benthamiana* leaves, approximately 15 hours after previous infiltration with *A. tumefaciens* GV3101 carrying wild-type I2 or I2 mutants at a final OD600 of 0.3. Leaves were incubated for up to 7 days on wet paper towels in trays with 100% relative humidity. The trays were kept at room temperature. Quantification of the total area of infection was carried out using the Image J software (Schneider, Rasband et al. 2012). Mycelial growth of *P. infestans* 88069td strain was visualized using a Leica Stereomicroscope (Leica Microsystems CMS GmbH) mounted with a CCD camera under UV LED illumination and filter settings for DsRed. In this case, the lesion area was monitored using bioimage analysis software. The software plugin was designed so that it could batch process a series of TIFF microscope images based on a number of core ImageJ/FIJI libraries. The algorithm extracted the intensity plane from the TIFF images, based on which masking methods were applied to identify regions with high intensity and contrast values. Tailored feature selection functions were implemented to detect objects such as the scale (pixel to μm) and infection areas on every image. Finally, a 2D convex hull method was applied to measure recognized infection areas and score the size/perimeter of the infection (in both pixel and μm).

### Western blots

Wild-type and mutant I2 and R3a proteins were transiently expressed by *A. tumefaciens*-mediated transformation in *N. benthamiana* leaves, and samples were collected at 2, 3 and 5 dpi. Protein extracts were prepared by grinding leaf samples in liquid nitrogen and extracting 1 g of tissue in 3 mL of GTEN protein extraction buffer (150 mM Tris-HCl, pH 7.5, 150 mM NaCl, 10% (w/v) glycerol, and 10 mM EDTA) with freshly added 10 mM dithiothreitol, 2% (w/v) polyvinylpolypyrrolidone, 1% (v/v) protease inhibitor cocktail (Sigma), and 1% (v/v) Nonidet P-40 as previously described (Win, Kamoun et al. 2011). Immunoblotting was performed with a polyclonal antibody raised against the CC domain of R3a (a-R3a).

### I2 structure predictions

Homology models of the individual CC and NB-ARC regions of I2 were generated using protein fold recognition algorithms, as implemented by 4 different servers Intfold2 (Buenavista, Roche et al. 2012, McGuffin, Atkins et al. 2015), iTASSER (Zhang 2008), Phyre2 (Kelley, Mezulis et al. 2015) and SWISS-MODEL (Biasini, Bienert et al. 2014). For the CC domain, three of the prediction programs (iTASSER, Phyre2 and SWISS-MODEL) identified the CC domain of the barley CNL protein MLA10 (PDB code 3QFL) as the top-scoring template for modelling of the I2 CC domain. IntFOLD2 (Buenavista, Roche et al. 2012, McGuffin, Atkins et al. 2015) produced a model based on the CC domain of the potato CNL protein Rx (PDB code 4M70) which adopts a distinct structure, quite different at the three-dimensional level to MLA10. The NB-ARC domain was modeled using Apaf-1 (PDB code 1Z6T chain B) (Riedl, Li et al. 2005) as template, which was the top score of IntFold2. That template is in agreement with previously published experimental evidence showing that I2 has ATPase activity (Tameling, Elzinga et al. 2002), as determined by positioning of the key motifs known to be required for ATPases.

## Acknowledgements

This work was supported by the European Research Council (ERC, proposal “NGRB”), the UK Biotechnology and Biological Sciences Research Council (BBSRC, grants BB/I019557, BB/J00453), the John Innes Foundation and the Gatsby Charitable Foundation. AG received support from the Norwich Research Park fellowship and Onassis Foundation. The funders had no role in study design, data collection and analysis, decision to publish, or preparation of the manuscript. We thank Frank L. W Takken for providing the *I2* and *AVR2* clones. We also thank V.G.A.A.Vleeshouwers for providing the NL00228 strain and Steve Whisson for providing the 88069td strain.

## Footnotes

1 GenBank number KR108299

## Supplementary material

**Figure S1:**
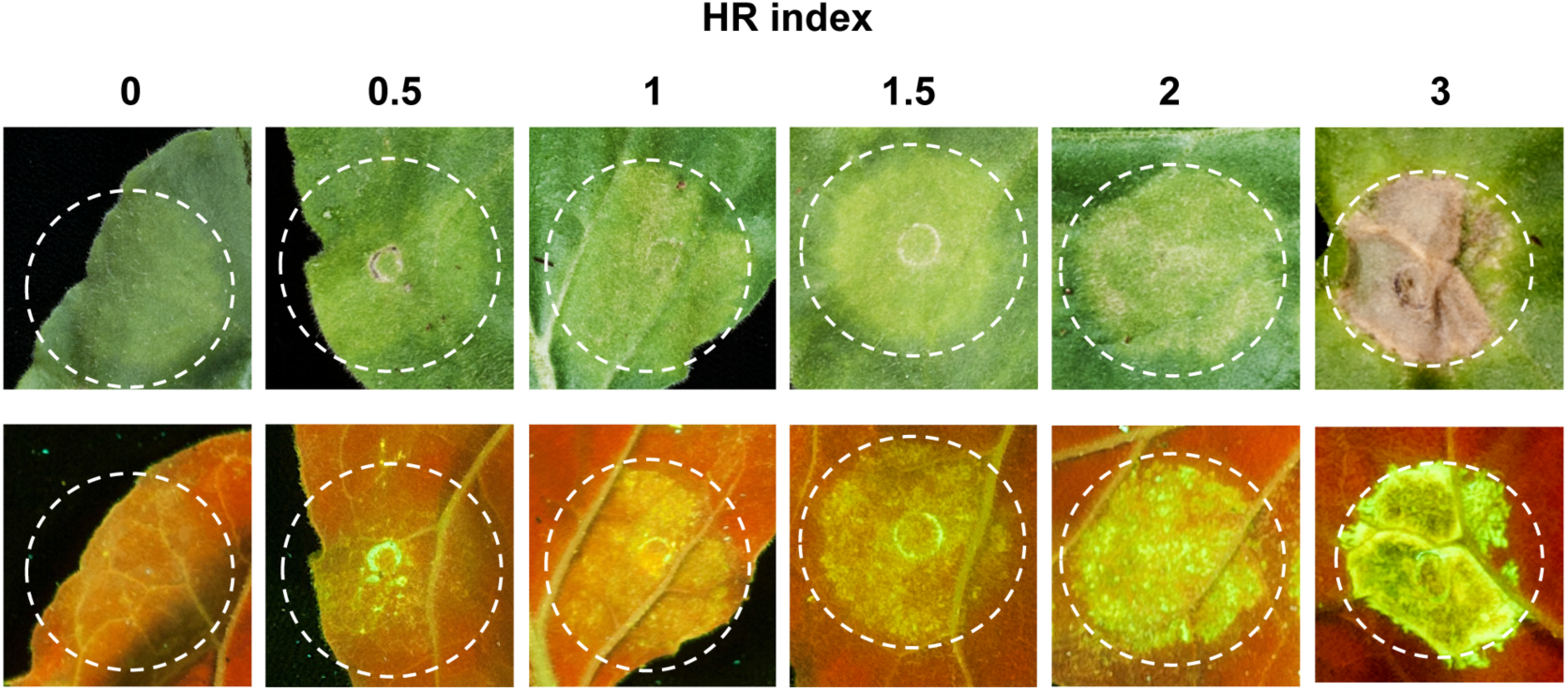
The hypersensitive response (HR) index. The index was scored according to an arbitrary scale from 0 (no HR phenotype) to 3 (confluent necrosis on the infiltrated area). Representative pictures for different values of HR index are shown.

**Figure S2:**
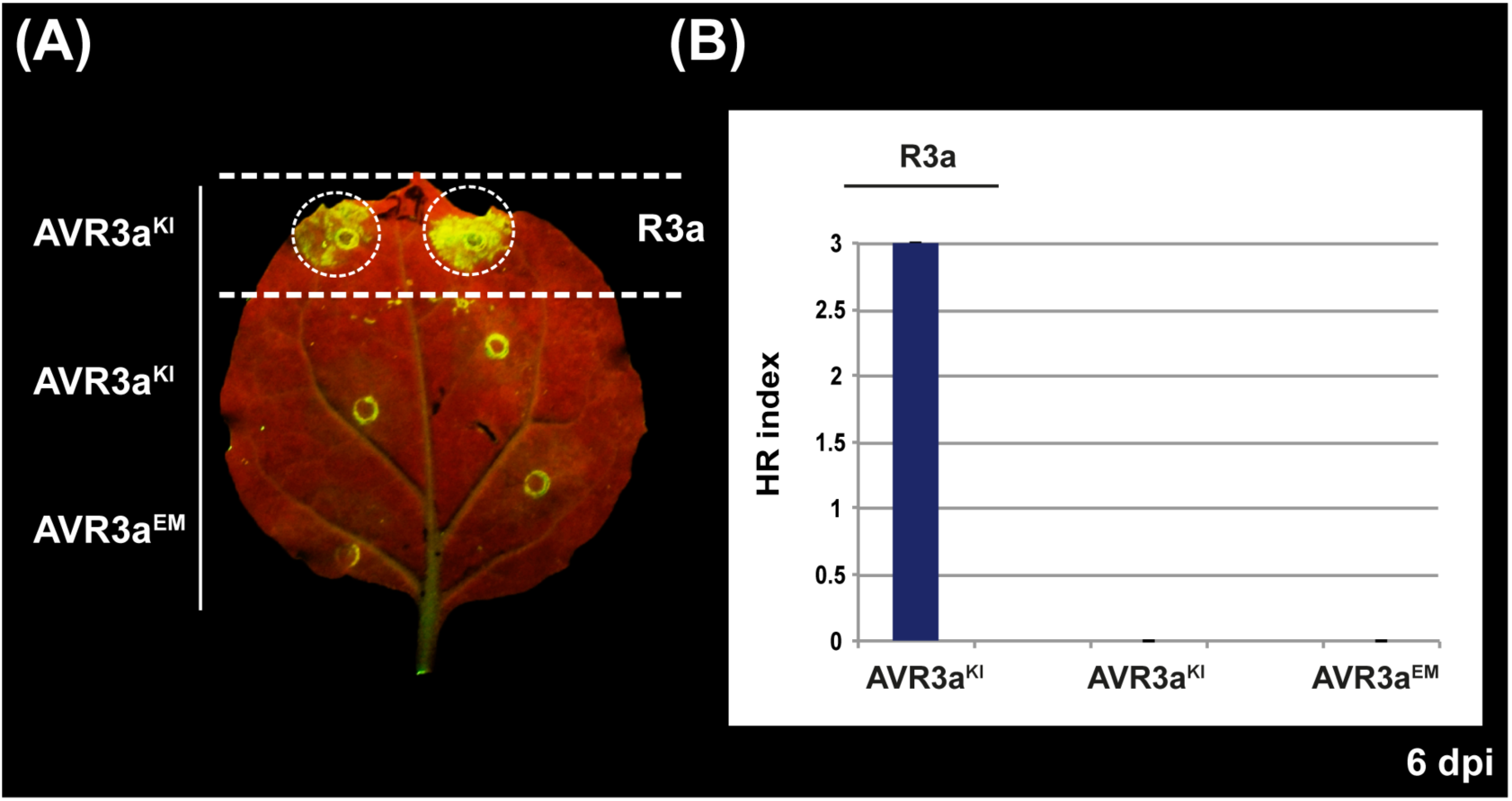
AVR3a variants do not cause cell death when after transient expression in *Nicotiana benthamiana*. A, Hypersensitive response (HR) phenotypes of the *Phythophthora infestans* AVR3a variants after transient expression in *N. benthamiana* leaves. Co–expression of wild-type R3a with the AVR3a^KI^ variant in *N. benthamiana* leaves was used as a positive control in this experiment. Wild-type *R3a* was under transcriptional control of the *Rpi-vnt1.1* promoter and AVR3a^KI^, AVR3a^EM^ were expressed from a *Potato Virus X* (PVX)-based vector. The picture was taken at 6 days post-infiltration (dpi). B, HR indices corresponding to the experiment described in A. Values scored at 6 dpi are plotted. Bars represent the average of 16 replicas for each combination of constructs; error bars represent standard deviation.

**Figure S3:**
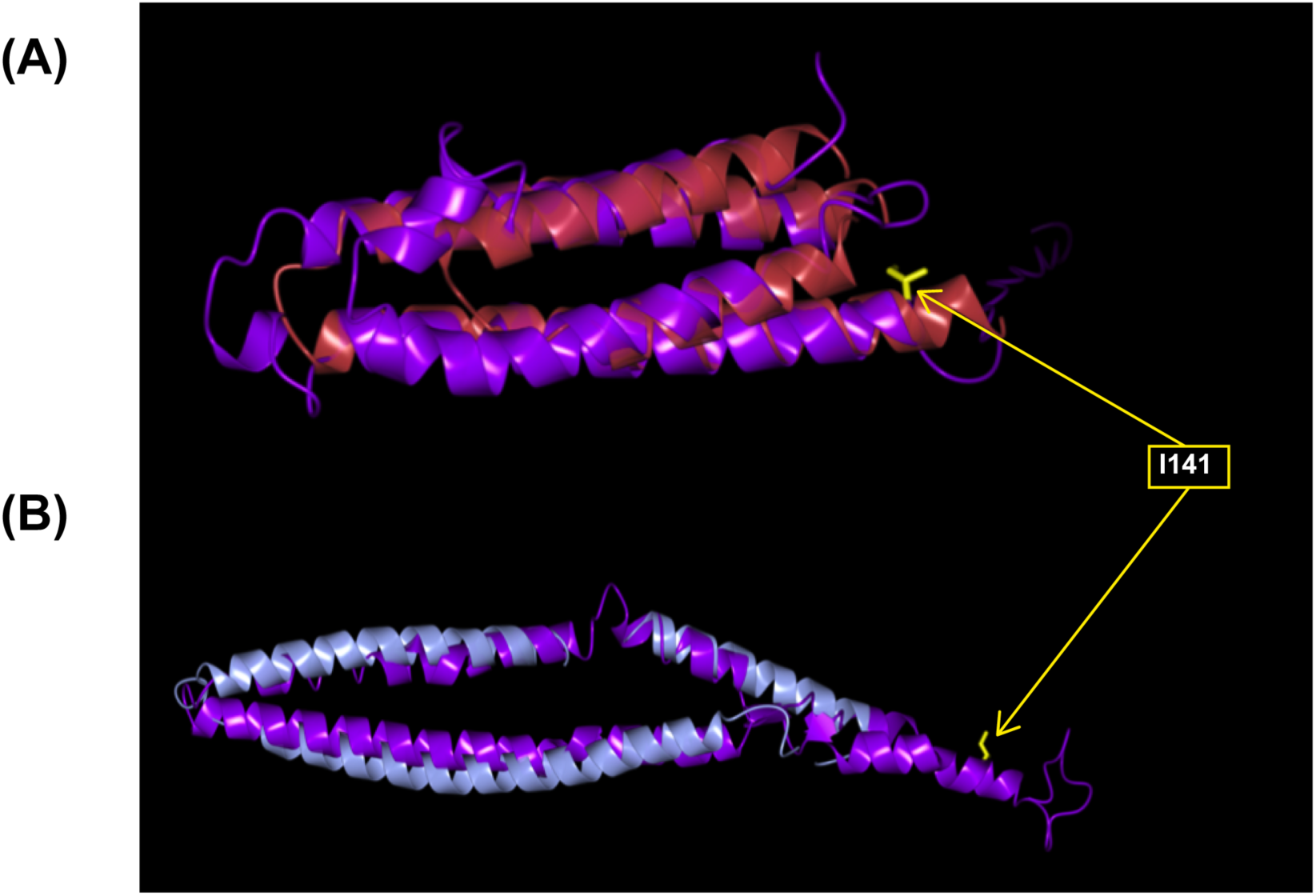
Modelling of the coiled-coil domain of I2 positions residue I141 C-terminal of defined coiled-coil region. Unbiased modelling of the CC domain of I2 resulted in the selection of the potato CNL protein Rx (blue) (A) and the barley CNL protein MLA10 (red) (B) CC domains as the templates giving the highest-confidence models. Homology models of the CC domain of I2 generated by IntFOLD2 (A) and iTASSER (B) are indicated in purple and are superposed onto the template structures with an rmsd of 3.25Å and 1.75Å respectively. Positions I141 are shown in yellow stick representation.

**Figure S4:**
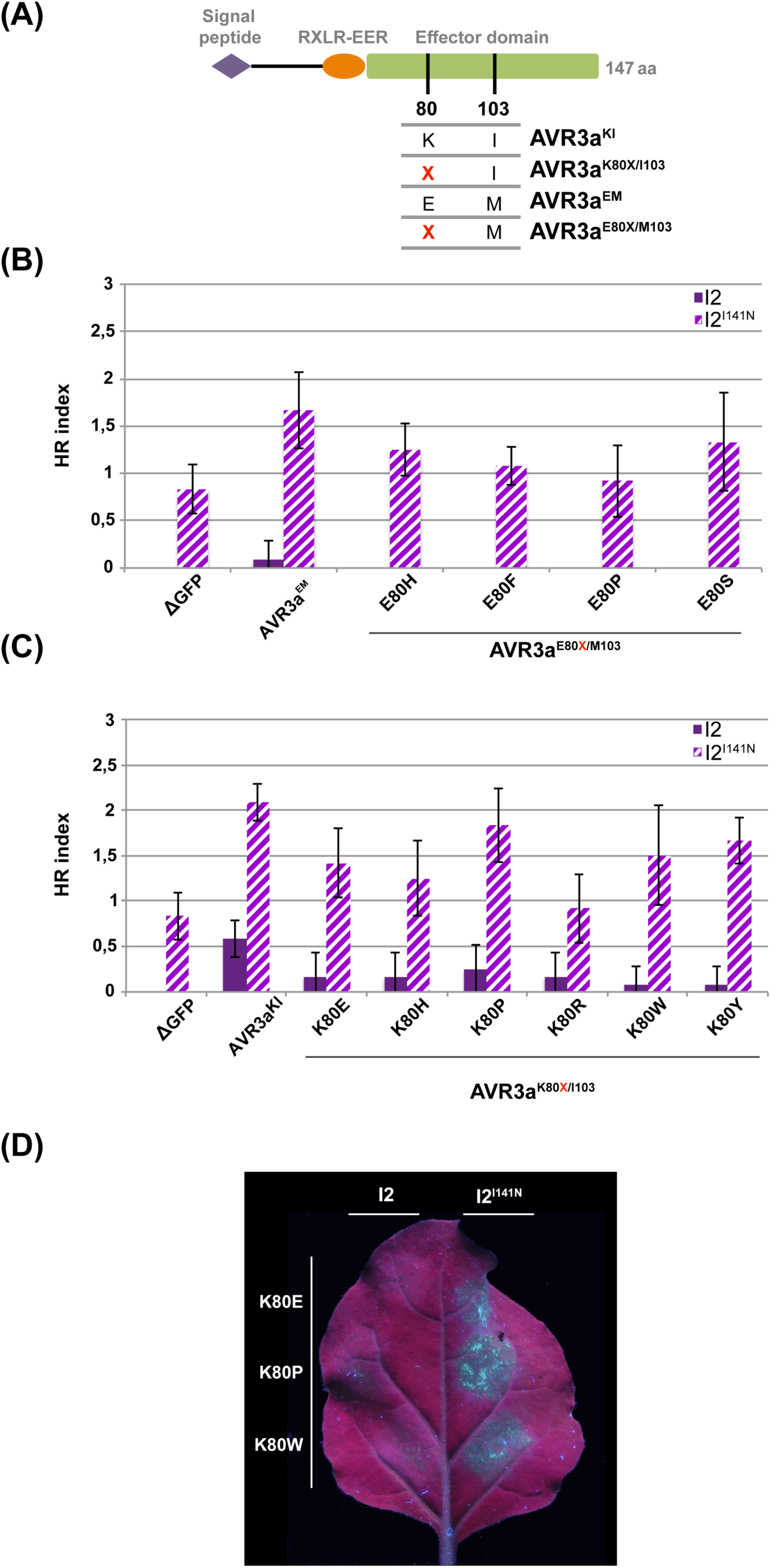
The I2^I141N^ mutant responds to multiple AVR3a variants. A, Schematic representation of *Phytophthora infestans* effector protein AVR3a, showing polymorphic amino acid positions 80 and 103 in the effector domain. B and C, Hypersensitive response (HR) indices of wild-type I2 and I2^I141N^ after co-expression with the *P. infestans* AVR3a variants (AVR3a^E80X/M103^ and AVR3a^K80X/I103^ in B and C respectively) in *Nicotiana benthamiana* leaves. The wild-type and mutant *I2* were under transcriptional control of the *Cauliflower mosaic virus* 35S promoter. A truncated version of green fluorescent protein (ΔGFP), AVR3a^KI^, AVR3a^EM^, and a collection of AVR3a mutants with amino-acid substitutions (as indicated) at position 80 in AVR3a^K80X/I103^ and AVR3a^E80X/M103^ were expressed from a *Potato virus X* (PVX)-based vector. Values scored 6 days post-infiltration (dpi) are plotted; columns represent the average of 6 replicas and error bars represent standard deviation. The experiment was performed several times with similar results. D, Representative picture showing differential HR after co-expression of wild-type I2 or I2^I141N^ with three AVR3a variants: AVR3a^K80E/I103^, AVR3a^K80P/I103^ or AVR3a^K80W/I103^. The picture was taken at 6 dpi.

**Figure S5:**
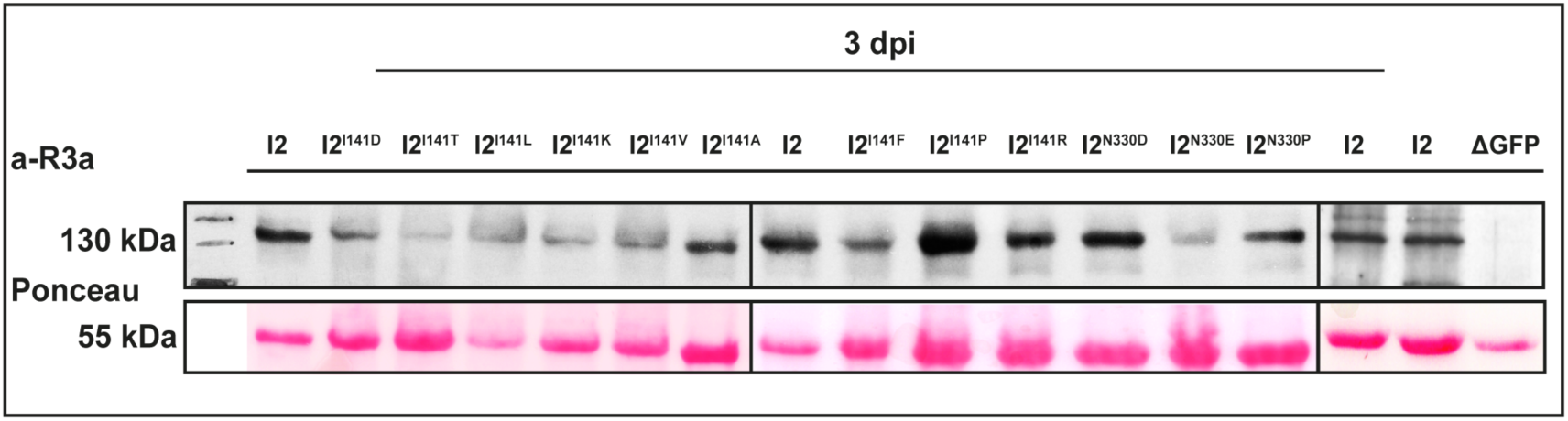
The I2 loss-of-response mutants are all expressed *in planta*. *Nicotiana benthamiana* leaves were agroinfiltrated with constructs to express wild-type and mutant I2 proteins or a truncated version of green fluorescent protein (ΔGFP). Total protein extracts from leaves sampled at 3 days post-infiltration (dpi) were subjected to SDS-PAGE followed by Immunoblotting with a polyclonal antibody raised against the CC domain of R3a (a-R3a, upper panel). A band of approximately 145 kDa corresponding to I2 was present in total extracts of the leaves infiltrated with the *I2* wild-type and mutant constructs. Ponceau S staining of Rubisco (lower panel) is shown as control of the amount of protein loaded and transferred in each lane. Sizes (in kDa) are indicated on the left. The different boxes indicate different blots.

**Figure S6:**
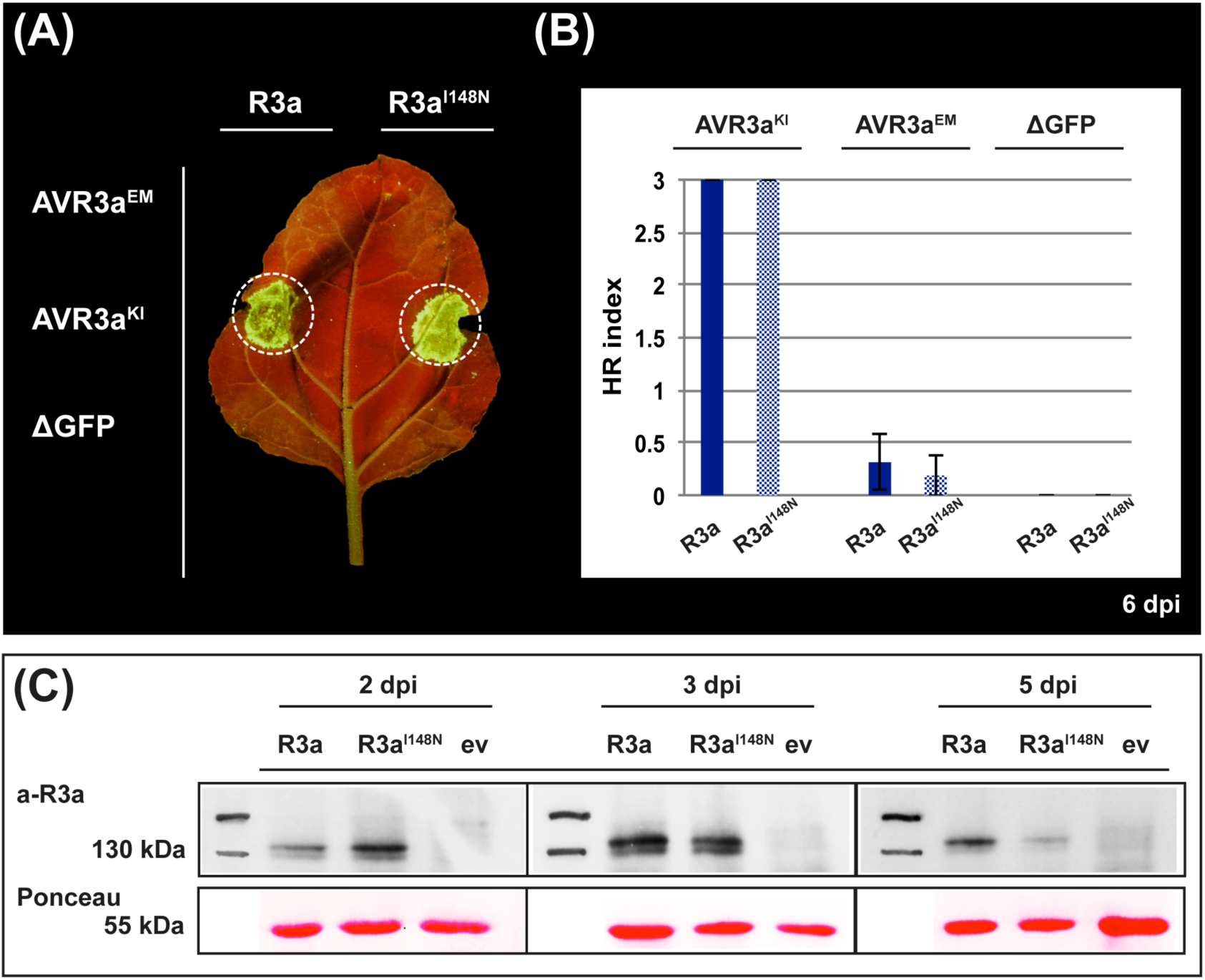
R3a^I148N^ response to the AVR3a variants is similar to that of wild-type R3a. A, Hypersensitive response (HR) phenotypes of wild-type R3a and R3a^I148N^ after co-expression with the *Phythophthora infestans* AVR3a variants in *Nicotiana benthamiana* leaves. Wild-type and mutant *R3a* were under transcriptional control of the *Rpi-vnt1.1* promoter and AVR3a^KI^, AVR3a^EM^ or a truncated version of green fluorescent protein (ΔGFP) were expressed from a *Potato Virus X* (PVX)-based vector. The picture was taken at 6 days post-infiltration (dpi). B, HR indices corresponding to the experiment described in A. Values scored at 6 dpi are plotted. Bars represent the average of 18 replicas for each combination of constructs; error bars represent standard deviation. C, Western blot assays were performed on total protein extracts from *N. benthamiana* leaves infiltrated with the *R3a* constructs described in A. The binary vector pCBNptII-vnt1.1P/T (Eugenia Segretin, Pais et al. 2014) was included as negative control (empty vector, ev). Total protein extracts were obtained at 2, 3 and 5 dpi and subjected to SDS-PAGE followed by immunoblotting with a polyclonal antibody raised against the CC domain of R3a (a-R3a). A band of approximately 146 kDa corresponding to R3a was present in the total extracts from the leaves infiltrated with the *R3a* constructs. Ponceau S staining of Rubisco (lower panel) is shown as control of the amount of protein loaded and transferred in each lane. Sizes (in kDa) are indicated on the left. The different boxes indicate different blots.

**Figure S7:**
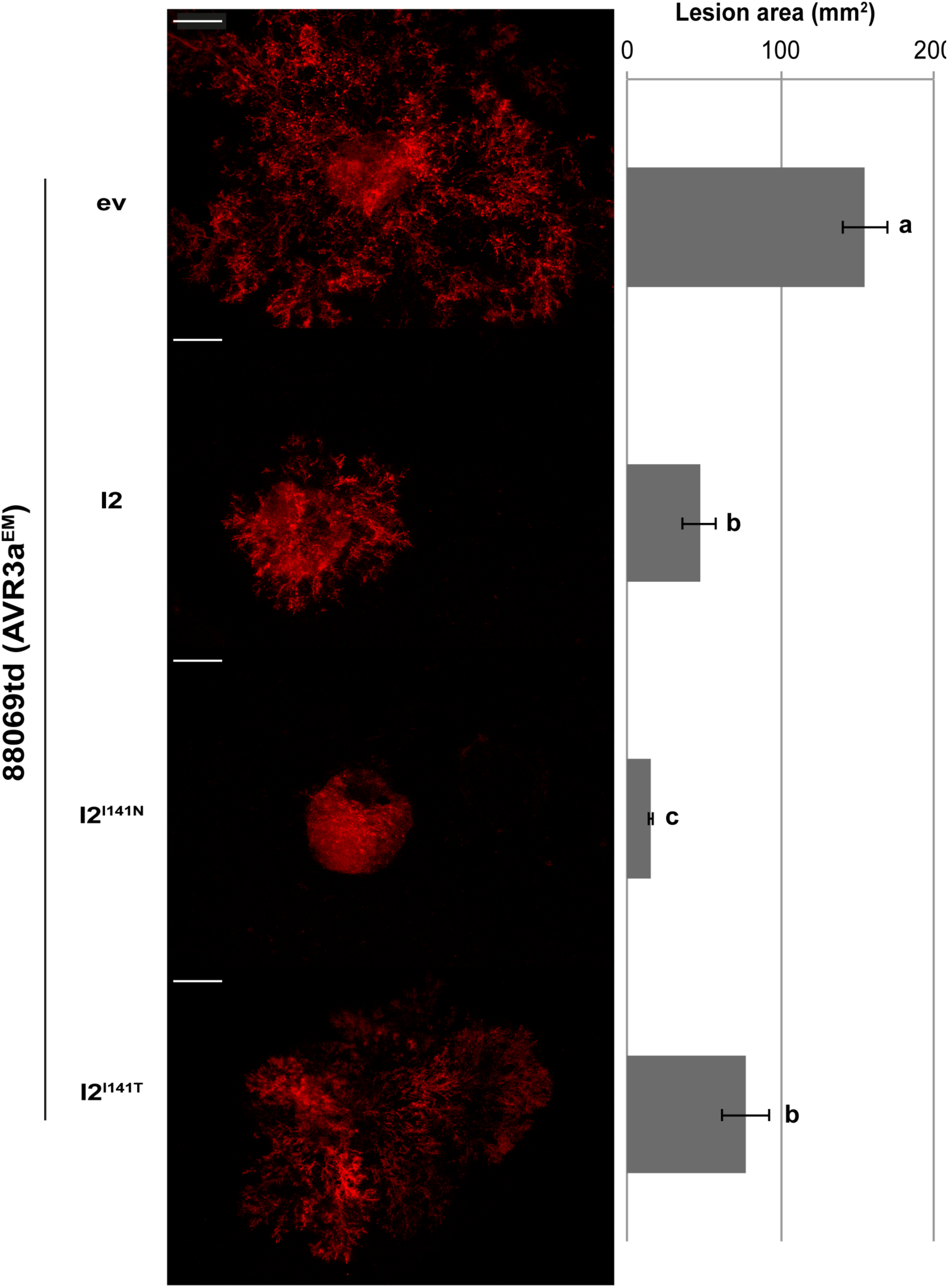
I2^I141N^ prevents *Phytophthora infestans* 88069td growth. Wild-type and mutant *I2* were expressed in *Nicotiana benthamiana* leaves under transcriptional control of the *Cauliflower mosaic virus* 35S promoter. After approximately 15 hours, the infiltrated leaves were drop-inoculated with a *P. infestans-*zoospore suspension corresponding to the 88069td strain (AVR3a^EM^). 88069td is a transgenic strain expressing the red fluorescent marker tandem-dimer RFP, known as tdTomato. The pictures were taken at 6 days post-inoculation. The area of *P. infestans* lesions was determined by analyzing the fluorescent signal with a bioimage analysis software (described in materials and methods). Values corresponding to 6 days post-inoculation are plotted. Bars represent the average of 24 replicas for each treatment; error bars represent standard deviation. The experiment was performed 3 times and a representative repeat is shown. The Gateway binary vector pK7WG2 was included as negative control (empty vector, ev). I2^I141T^ is a loss-of-response I2 mutant and was used as an additional negative control in this experiment. Significant differences between the groups are indicated by letters and were determined in an analysis of variance (ANOVA) (p≤10^-4^).

**Supplementary table S1:**
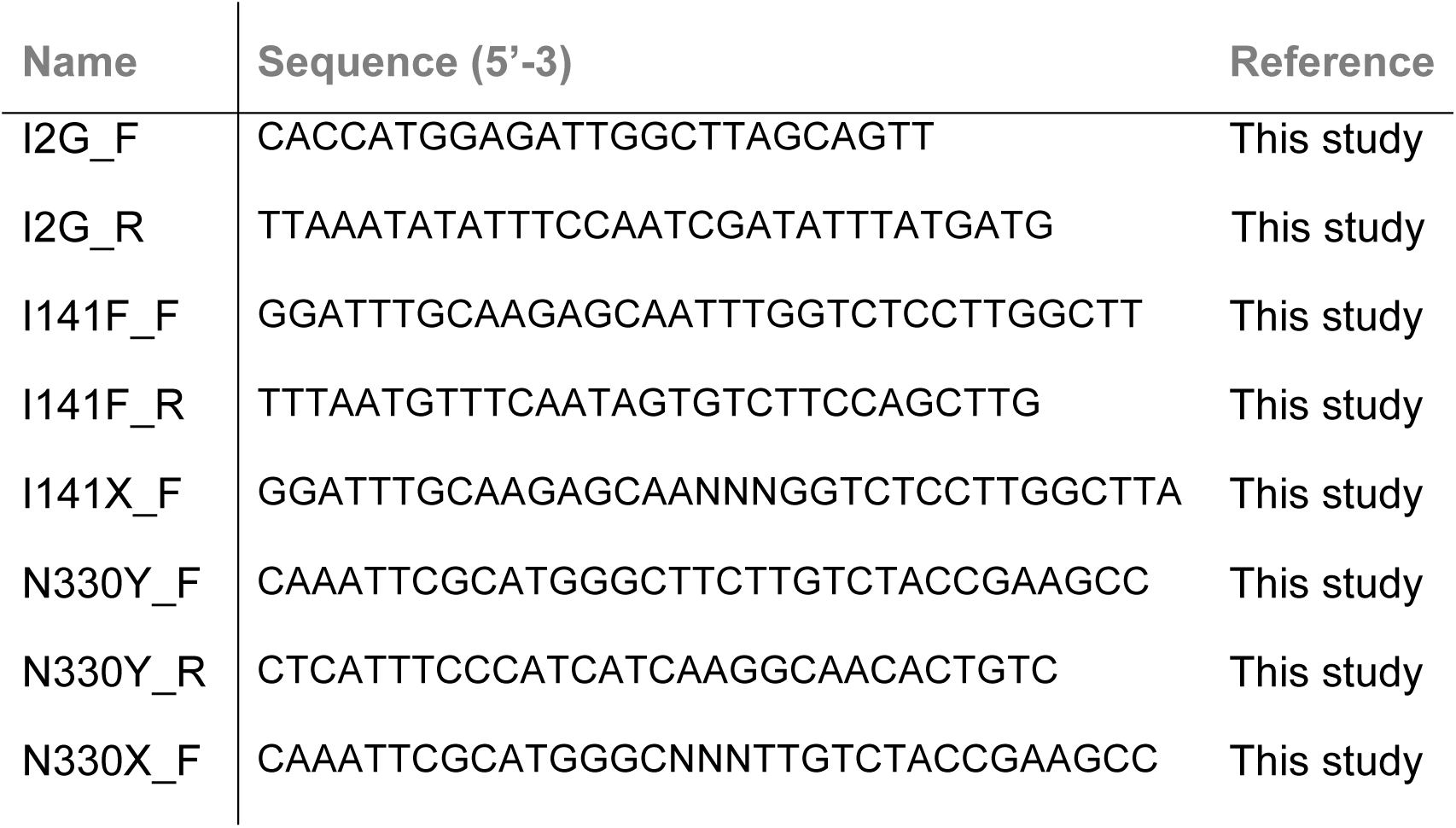
Primers used in this study.

